# Transcriptomic and lipidomic profiles reveal higher thermal adaptability in a tropical versus a temperate isolate of *Chromera velia*

**DOI:** 10.64898/2026.10.21.760691

**Authors:** Ayush Sharma, Aleš Tomčala, Tomáš Skalický, Dorsaf Ennaceur, Aleš Horák, Jitka Richtová, Vijaya Geetha Gonepogu, Jan Šlapeta, Miroslav Oborník

## Abstract

The apicomplexan alga *Chromera velia* is the closest known phototrophic relative of apicomplexan parasites (Sporozoa). Despite its scientific importance as a model for the evolution of parasitic Apicomplexa, only two strains are available in culture. The first isolate (SH), obtained from stony coral in the cold waters of Sydney Harbour, was used for the original species description. The second isolate (OTI) originates from tropical corals at One Tree Island in the Great Barrier Reef. In this study, we compare their growth, transcriptomic, and lipidomic profiles at 20°C and 26°C. Both strains grew faster at 26°C, yet they responded differently to higher temperature. In SH, warming downregulated nearly 85% of its thermally responsive genes, with motility-, cytoskeleton-, and lipid-associated genes most strongly affected, while at 26°C SH retained higher expression of translation and nitrogen, amide, and peptide biosynthesis genes than OTI. OTI, by contrast, mounted a smaller, more balanced response and maintained stable expression of cilium, axoneme, and microtubule genes across both temperatures. The lipidomes of the two strains showed both shared and divergent responses. Both isolates sense and respond to increasing temperature in the same basic way, but OTI stabilizes its membranes, whereas SH redirects lipid synthesis from membrane lipids to storage lipids. OTI membranes were also constitutively less unsaturated than those of SH at both temperatures, consistent with a membrane already suited to warmer water that needs less remodeling. Our data show that two isolates of the same species, originating from different thermal environments, respond to elevated temperature through distinct transcriptomic and lipidomic programs. This suggests that the tropical OTI isolate is adapted to higher temperatures and demonstrates how reef-associated algae respond to rising ocean temperatures.

## Introduction

Apicomplexa is a group of alveolate protists that possess a complex plastid [1]. The group consists of two branches: Sporozoa, which are medically highly significant parasites of animals, including humans [2], and Apicomonada [3] a trophically diverse group of protists that includes phototrophs (chromerids), predators (colpodellids), and parasites (*Piridium sociabile*) [4,5]. Almost all apicomplexans contain a plastid: either a photosynthetic plastid (chromerids), a non-photosynthetic plastid with a genome (most sporozoans and *P. sociabile*), a non-photosynthetic plastid without a genome (colpodellids and some sporozoans such as gregarines), or the plastid organelle may have been lost entirely, as in the sporozoan genus *Cryptosporidium* and some gregarines [5–7]. Phylogenetic analysis clearly shows that photosynthesis has been lost multiple times within Apicomplexa [8], but it appears that all apicomplexan plastids share a single, originally photosynthetic origin [9].

Chromerids (Apicomplexa, Apicomonada) are a polyphyletic group of apicomplexan algae isolated from stony corals in Australia [10]. Their phylogenetic position within Apicomonada (also called chromopodelids) classifies chromerids as the closest known photosynthetic relatives of apicomplexan parasites (Sporozoa) [10,11]. Only two species of chromerids have been described: *Chromera velia* [10,11] and *Vitrella brassicaformis* [12]. While *V. brassicaformis* is very likely an epiphytic alga growing on the coral surface [13], *C. velia* is probably a photoparasite of coral larvae [5,14,15]; however, it has never been found in adult corals. The two chromerids exhibit very different morphology and ultrastructure but have similar pigmentation, as they lack chlorophyll *c*, the hallmark of algae with rhodophyte-derived complex plastids [11]. *Vitrella brassicaformis* has a complex life cycle that includes sexual reproduction [16], while the life cycle of *C. velia* is reduced and likely asexual [11].

Chromerids appear to be abundant worldwide according to environmental sequencing [17]. Recently, metagenomic data have shown great diversity of chromerids associated with microbialites in marine and freshwater habitats across the globe [18]. However, none of these chromerids is available in culture. Only one isolate, described as *C. velia*, is available in public culture collections. This isolate (SH; CCMP 2878 and CCAP 1602) was obtained at Sydney Harbour (Fig 1A) from the scleractinian coral *Plesiastrea versipora* [10]. The second isolate (OTI) used in this study, was obtained from the scleractinian coral *Leptastrea purpurea* at One Tree Island in the Great Barrier Reef (GBR) (Fig 1A) [19,20]. This strain was referred to as *Chromera velia* by previous authors [19,20]. Here we refer to it as *C. velia* (OTI strain) throughout this manuscript. Three other isolates have been mentioned in the literature: *C. velia*, isolated from stony coral *Montipora digitata* in the central part of the GBR [14,21], see S1 Table), *Chromera* sp., isolated in the Caribbean Sea from the lettuce coral *Agaricia agaricites*, whose culture has unfortunately been lost [22], and *Chromera* velia (GBR strain) isolated from GBR by Mohamed and group [15]. The two isolates in our lab originate from different thermal environments: the Average water temperature at Sydney Harbour is about 20 °C (SH), while at One Tree Island it is 26 °C (OTI).

**Figure 1.**
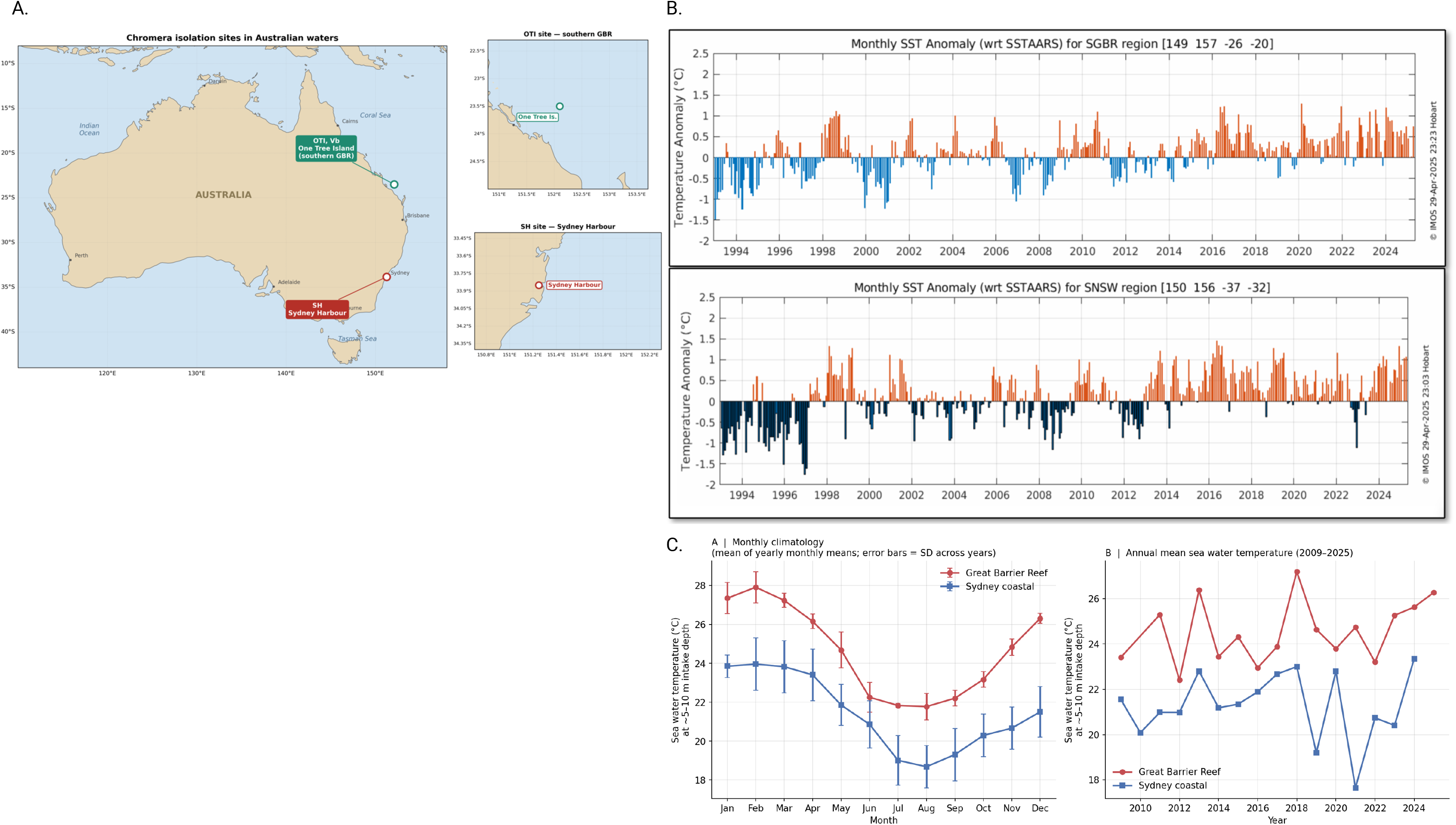
Geographic origin and thermal regimes of the two *Chromera velia* strains used in this study. **(A)** Map of Australia showing the two coastal isolation sites, with detailed insets (right). The OTI strain (green) was isolated from One Tree Island, southern Great Barrier Reef (23.50°S, 152.10°E; Queensland); the SH strain (red) from Sydney Harbour (33.84°S, 151.28°E; New South Wales). **(B)** Monthly sea-surface temperature (SST) anomalies, 1993–2025, relative to _C._ the long-term satellite-derived SSTAARS climatology, for bounding boxes covering the southern Great Barrier Reef (SGBR; 149–157°E, 26–20°S; top) and southern New South Wales (SNSW; 150–156°E, 37–32°S; bottom), corresponding to the OTI and SH isolation sites respectively. Warm anomalies are shown in orange and cool anomalies in blue. **(C)** Monthly climatology, calculated as the mean of yearly monthly mean sea-water temperatures, for the Great Barrier Reef (red; OTI site) and Sydney coastal (blue; SH site) regions; error bars denote standard deviation across years. *Right:* Annual mean sea water temperature for the same two regions from 2009 to 2025. Only quality-controlled observations (QC flag = Z) within each regional bounding box were included. Data in (B) and (C) accessed from the IMOS OceanCurrent portal (https://oceancurrent.aodn.org.au).

Because we have preferentially used the formally described SH strain [10] in our experiments, and because 26°C has generally been regarded as a favorable temperature at which *C. velia* grows faster, we have historically used it as our standard cultivation condition. Given this, and given that *C. velia* appears globally abundant in environmental sequencing surveys [17,18], we set out to test whether the origin of the SH and OTI isolates shapes their physiology and gene expression, or whether they behave uniformly regardless of origin. To address this, we compared the growth, transcriptomic and lipidomic profiles of both isolates at 20°C and 26°C, temperatures that bracket the thermal conditions typical of their two sites of origin (Fig 1). Both strains grew faster at 26°C than at 20°C, confirming it as a favorable growth temperature. At the molecular level, however, the strains responded to the same temperature increase in markedly different ways: SH showed a transcriptional response dominated by downregulation, whereas OTI’s response was smaller and more balanced. In the lipidome, both isolates shared part of their response to increased temperature but differed in how they rebuilt their membranes, and OTI membranes were less unsaturated than those of SH at both temperatures. Despite the lack of morphological divergence, the two isolates thus differ substantially in how they respond to the same thermal conditions.

## Materials and Methods

### Cultures and cultivation

*Chromera* velia strain (SH) [10] was obtained from the Culture Collection of Marine Phytoplankton (CCMP) (Boothbay Harbor, Maine). *Chromera velia* strain (OTI) was kindly provided by Jan Šlapeta from the University of Sydney, Australia [19]. Both strains were grown in f/2 medium under 12/12 light/dark regime at 26 °C and 20 °C for prolonged period of times.

### Light and electron microscopy

Both coccoid cells and zoospores were observed for SH and OTI strains using an Olympus DP20 model BX53 microscope. For scanning electron microscopy, samples were prepared according to [11] and examined with a JEOL JSM 7401-F field emission scanning electron microscope. For transmission electron microscopy, cultured cells were pelleted by centrifugation at 15,000 rpm for 5 minutes and frozen using a Leica EM Pact high-pressure freezer. Freeze substitution was performed as described in [11], and samples were examined with a JEOL JEM 1010 transmission electron microscope equipped with a MegaView III digital camera (SIS).

### Climate data

To place strain-specific responses in a realistic thermal context, we compiled ocean temperature records for both isolation regions using two complementary metrics. Sea surface temperature (SST) refers to the temperature of the uppermost ocean skin (∼0–1 m) retrieved from satellite radiometers and is well suited to detecting long-term regional anomalies and marine heatwave events. In situ sea water temperature refers to the water immediately below the surface, measured directly by ship-mounted sensors, and better represents the depth range inhabited by *Chromera velia* and its coral hosts. Monthly SST anomalies from 1993 to 2025 were obtained from the IMOS OceanCurrent portal (https://oceancurrent.aodn.org.au), expressed relative to the SST Atlas of Australian Regional Seas (SSTAARS) climatology, for bounding boxes covering the southern Great Barrier Reef (SGBR; 149–157°E, 26–20°S) and southern New South Wales (SNSW; 150–156°E, 37–32°S), corresponding to the OTI and SH isolation sites.

Absolute *in situ* sea water temperatures at ∼5–10 m depth were obtained from the IMOS Ships of Opportunity Sea Surface Temperature Sub-Facility (SOOP-SST) via the Australian Ocean Data Network portal (https://portal.aodn.org.au); despite the “SST” label in the dataset name, these measurements come from ship engine cooling-water intakes and represent near-surface water rather than the ocean skin. Records from 2009 to 2025 were restricted to quality-controlled observations (QC flag = Z) within regional bounding boxes for the southern GBR (10–24°S, 142–154°E; OTI site) and Sydney coastal region (33.5–34.0°S, 151.2–152.3°E; SH site). Monthly climatology was calculated as the mean of yearly monthly means, and annual means as the mean of all valid observations per year. Climate raw data can be accessed through our online repository.

### Growth of *Chromera* strains at 20 °C and 26 °C

Both SH and OTI strains were analyzed for growth under two temperature conditions, following the protocol recently published by Richtová et al[23] for the culture and analysis of *Chromera velia* isolates. Cultures were maintained in sterile T75 Erlenmeyer flasks containing 50 mL of f/2 medium under a 12:12 hour light:dark photoperiod at 40 μmol photons m□² s□¹, with growth experiments conducted at 20 °C and 26 °C in temperature-controlled incubators (±0.5 °C accuracy). Each strain was cultured in biological quadruplicate at each temperature (n = 4), resulting in a total of 32 experimental flasks (16 for growth monitoring, 16 for zoospore quantification). Cultures were inoculated at an initial optical density of approximately 0.075– 0.08 (OD □) and monitored every 3 days over 15 days. Growth was quantified by measuring optical density at 600 nm with a spectrophotometer, using samples transferred to 96-well plates. Specific growth rate (μ, day ¹) was calculated from Day 0 and Day 15 OD values as μ = [ln(OD □□□) − ln(OD □)] / 15. Zoospore numbers were counted daily for 9 days by direct microscopic examination within the culture flask, using four technical replicate fields per flask across four biological replicate flasks per condition (n = 4 flasks, 16 fields total per condition) [23].

### RNA Isolation

*C. velia* strains OTI and SH grown at 20 °C and 26 °C were harvested in logarithmic phase (day 7) onto 0.22 µm cellulose acetate filters (380–500 mg biomass per replicate), rinsed with RNAlater, flash-frozen in liquid nitrogen, and stored at −80 °C. All solutions were DEPC-treated where possible. Frozen filters were ground to a fine powder under liquid nitrogen and lysed in 3 mL ice-cold TRI Reagent RT (Molecular Research Center, cat. no. RT 111) with 1% β-mercaptoethanol, following the manufacturer’s protocol with two sequential chloroform extractions. RNA was further purified on a QIAcube Connect using the RNeasy Mini Kit (Qiagen, cat. no. 74104) with on-column DNase digestion, precipitated overnight with sodium acetate/isopropanol, resuspended in nuclease-free water, and stored in GenTegra RNA tubes (cat. no. GTR5025-S). Purity was assessed by NanoDrop (mean A260/280 = 2.13 ± 0.02; A260/230 = 2.42 ± 0.69; n = 16 extractions, of which 12 were sequenced) and RNA integrity was checked using Agilent 2100 Bioanalyzer (Agilent Technologies).

### Sequencing of transcriptomes

Three independent biological replicate transcriptome libraries of SH and OTI isolates were prepared from cultures grown at 20°C and 26°C using the TruSeq Stranded mRNA Library Prep Kit, and sequenced on the Illumina NovaSeq 6000 platform (2x150 bp paired-end reads, minimum 50 million reads per sample). The corresponding raw reads are available in GenBank under accession numbers PRJNA1519432. Adapters were trimmed from the raw reads using Trimmomatic v. 0.39 [24], and data quality was assessed with FastQC v. 0.12. Reads were mapped to the CryptoDB-54 *C. velia* CCMP2878 genome reference using STAR v. 2.7.10b [25]. To define the expressed (core and pan) transcriptome, raw counts were converted to counts per million (CPM) using the cpm() function of edgeR v3.28.0[26] (edgeR was used only for CPM calculation, not for differential expression testing); a gene was considered expressed in a condition if it reached CPM >= 1 in all three biological replicates. Differential expression between strains and between temperatures was tested with DESeq2 [27], using a low-count pre-filter (>= 10 counts in at least three samples); genes with a false discovery rate (FDR, Benjamini-Hochberg adjusted p) < 0.05 and an absolute log2 fold change >1 were considered significantly differentially expressed and used for downstream analysis. The raw differential expression analysis (DEA) tables, provided as Excel sheets, are available through our online repository

### Functional annotation and GO enrichment analysis

Gene annotations and GO enrichment analyses were performed using CryptoDB (VEuPathDB) release 54 [28], which hosts the *Chromera velia* CCMP2878 genome. Product descriptions, gene types and genomic coordinates for differentially expressed genes were retrieved by uploading gene ID lists to the CryptoDB gene record search and exporting the resulting gene tables. For enrichment analysis, DEG lists were partitioned by contrast and direction and submitted to the CryptoDB GO Enrichment tool, using the *C. velia* CCMP2878 whole genome as background. Analyses used the full ontology rather than the GO Slim subset, included both computed and curated evidence codes, and applied a P-value cutoff of 0.05. Enrichment was run separately for the three GO ontologies: Biological Process, Cellular Component and Molecular Function. For each enriched term the tool returns background and result gene counts, fold enrichment, odds ratio and raw P-value, together with Benjamini-Hochberg and Bonferroni adjusted values. Enrichment was performed for two groups of gene sets. For the strain comparison, we analysed genes more highly expressed in OTI at 26 °C (OTI heat-up, n = 1,090), in OTI at 20 °C (OTI cool-up, n = 311), in SH at 26 °C (SH heat-up, n = 510), and in SH at 20 °C (SH cool-up, n = 811), together with the two direction-flip sets, in which the strain difference reverses between temperatures (higher in OTI at 20 °C and in SH at 26 °C, n = 4; higher in OTI at 26 °C and in SH at 20 °C, n = 69). For the thermal response, we analysed genes uniquely upregulated within each strain at each temperature (OTI-unique, up at 20 °C n = 1,155 and up at 26 °C n = 1,320; SH-unique, up at 20 °C n = 2,450 and up at 26 °C n = 447), together with the genes shared between strains: concordant (downregulated at 26 °C in both strains, n = 1,283; upregulated at 26 °C in both strains, n = 241) and discordant (downregulated in SH and upregulated in OTI, n = 86; upregulated in SH and downregulated in OTI, n = 4). Terms were considered significantly enriched at Benjamini-Hochberg FDR < 0.05. GO shows Biological Process results only. Each workbook contains three sheets (DEG, Annotation and Gene_ontology), and the Gene_ontology sheet holds the Biological Process, Cellular Component and Molecular Function tables. All Excel sheets are available through our online repository.

### Lipid extraction

*Chromera* cultures, four replicates per group, were harvested by centrifugation using a Micro 200 R Hettich centrifuge set to 20 °C for 10 min at 10,000 rpm. The supernatant was discarded, and the pellet was used for lipid extraction following the protocol from [29] with slight modifications. The pellet was briefly homogenized in 0.7□mL of chloroform:methanol (2:1, v/v) in BeatBUG® vials using a BeatBUG® homogenizer (two 10□s cycles at 400 rpm). After homogenization, the mixture was centrifuged at 4□°C for 5□min at 10,000□rpm, and the resulting supernatant was transferred to a 2 mL Eppendorf tube. The homogenization step was repeated for the remaining pellet. Subsequently, 0.4 mL of deionized water was added to the combined supernatants, which were vortexed vigorously, and centrifuged again under the same conditions (4 °C, 5 minutes, and 10,000 rpm). After phase separation, the lower organic (lipid) phase was collected into reweighed tubes and evaporated under a stream of nitrogen. The final lipid content was determined gravimetrically using a Mettler Toledo XP6 Excellence Plus XP Micro Balance (6.1□g × 1□µg; Greifensee, Switzerland). Homogenates were dried and stored at −80 °C for further analysis.

### HPLC MS/MS

Two milligrams of crude fat extract from samples were dissolved in 500 µL of chloroform:methanol (1:2, v/v) containing the internal standard PC 17:0/17:0 (Sigma-Aldrich). Aliquots of each extract were analyzed for lipid composition by HPLC using an Accela liquid chromatograph with an autosampler (Thermo Fisher Scientific, San Jose, USA). Samples (5 µL) were injected onto a Gemini column (250□×□2□mm, 3□µm i.d.; Phenomenex, Torrance, USA). A linear ion trap LTQ-XL mass spectrometer (Thermo Fisher Scientific) was used in both positive and negative ESI modes. Instrumental parameters followed the methodology described in [30]. Data acquisition and processing were performed using Xcalibur software version 4.01 (Thermo Fisher Scientific, San Jose, CA, USA). Individual lipid species were identified based on their m/z values, retention times, ionization behavior in both polarity modes, and characteristic fragmentation patterns (for details, see [30]).

## Results

### Climate data

*Chromera velia* was isolated from *scleractinian* corals collected at geographically distant locations along the Australian coast (Fig 1A) at two sites: Sydney Harbor (SH; Wedding Cake West, 33.84°S, 152.28°E) in New South Wales and One Tree Island (OTI; 23.50°S, 152.10°E) on the southern GBR. Sydney Harbor and the GBR region are separated by approximately 10 degrees of latitude (∼1,100 km). To analyze the natural conditions of both isolates, we used seawater temperature data obtained from the Integrated Marine Observing System (IMOS) through the Australian Ocean Data Network (AODN) Portal. The data we used originate from the MARVL3 Australian shelf temperature data atlas, which compiles measurements from multiple sources, including IMOS, the Australian Institute of Marine Science (AIMS), CSIRO Ocean and Atmosphere, the Royal Australian Navy, and the World Oceans Database [31]. The temperature measurements were collected at depths of 0 - 5 m, corresponding to the shallow reef environments where the host corals of *C. velia* were collected.

Historical temperature data from 1993 to 2025 reveal distinct patterns of thermal anomalies in both regions (Fig 1B). The southern GBR region, including One Tree Island, showed a noticeable shift in Sea surface temperature (SST) patterns over the three-decade period (Fig 1B). From 1993 to about 2006, negative temperature anomalies predominated, with occasional warm periods. The most significant negative anomalies reached approximately −1.5 °C below the climatological average. After 2006, positive anomalies became more frequent. This trend is especially apparent after 2014, with intense warm anomalies, sometimes exceeding +1.0 °C above the average. Several distinct warming peaks are evident in 1998, 2016-2017, and 2020-2024, corresponding to known marine heatwave events and El Niño periods.

The southern New South Wales region (Sydney Harbour) showed a similar thermal pattern (Fig 1B), but with notable differences. During the early period (1993-2000), negative anomalies were observed, with strong cooling events in 1996-1998 reaching nearly −2.0 °C below average. The transition to more frequent positive anomalies occurred gradually, with an increase in warming events after 2014. The 2015-2020 period was characterized by persistent positive anomalies, indicating a sustained warming trend in Sydney coastal waters. Sydney Harbour experienced more extreme cold anomalies in the 1990s compared to the GBR region, but the recent warming trend has been consistent across both locations (Fig 1B). These long-term patterns indicate that both regions have shifted toward warmer conditions over the past three decades, with the most significant warming occurring in the last decade. These trends align with broader patterns of ocean warming in Australian waters documented by numerous studies [32,33], and the effects of these changes on local symbionts have also been studied [34].

### Monthly Temperature Profiles and Regional Differences in Sea water temperature

Analysis of monthly sea water temperatures reveals distinct patterns between the two study sites (Fig 1C). Both locations follow Southern Hemisphere seasonal patterns, with Sydney Harbour reaching peak temperatures in April (24.17°C) during late austral summer/early autumn, while the GBR peaks earlier in February (27.94°C) during austral summer. Both sites experience minimum temperatures in August during austral winter. The thermal profiles show that the GBR maintains warmer water temperatures than Sydney Harbour throughout the annual cycle, with an average difference of approximately 3.4°C (Fig 1C). This temperature difference reflects the latitudinal gradient between temperate Sydney Harbour (33.84°S) and the tropical GBR region (∼20°S). The robust dataset (186,243 temperature records for the Great Barrier Reef vs. 980 records for Sydney Harbour) provides climatological averages spanning 16 years of observations. The GBR region, where the OTI strain was isolated, consistently maintained sea water temperatures 1.4-4.8°C higher than those in Sydney Harbour throughout the year, with the maximum difference of 4.8°C in December and the minimum difference of 1.4°C in June (Fig 1C).

### Growth of *Chromera* strains at 20 °C and 26 °C

*Chromera velia* strains SH and OTI were grown at 20°C and 26°C, with optical density (OD □□□) monitored every three days from day 0 to day 24 (n = 4 biological replicates per condition) (Fig 2A). Both strains reached higher final OD at 26°C than at 20°C: by day 15, OD□□□ was 0.368 ± 0.026 (SH-26) and 0.371 ± 0.010 (OTI-26) at the higher temperature, compared with 0.204 ± 0.019 (SH-20) and 0.205 ± 0.007 (OTI-20) at the lower temperature. Specific growth rate (µ), calculated over the full day 0–15 interval, was higher at 26°C than at 20°C in both strains (SH: 0.103 ± 0.005 vs 0.065 ± 0.009; OTI: 0.102 ± 0.003 vs 0.067 ± 0.002). A two-way ANOVA (strain × temperature) revealed a significant effect of temperature (F□, □□ = 197.1, p = 8.3 × 10) but not strain (F□, □□= 0.08, p = 0.78) on specific growth rate, with no significant strain × temperature interaction (F□, □□= 0.32, p = 0.58) (Fig 2C). Independent two-sample t-tests confirmed the temperature effect within each strain (SH: t□ = 7.72, p = 2.5 × 10□□; OTI: t□ = 20.83, p = 8.0 × 10□□). Zoospore production was assessed daily from day 0 to day 9 in both strains at 20°C and 26°C (n = 4 biological replicates per condition). No zoospores were detected in either strain at 20°C at any timepoint. At 26°C, zoospores were absent through day 3 in both strains, then appeared transiently: SH-26 rose sharply from day 4 (mean ≈ 6/field) to a peak around day 6 (mean ≈ 32/field, with substantial variability across replicates), before declining to near zero by day 8–9. OTI-26 showed a much smaller and earlier peak, around day 4–5 (mean ≈ 2/field), and had returned to near zero by day 6 (Fig 2B). Zoospore counts at 26°C were therefore higher in SH than in OTI across the period of peak production, and this transient wave was specific to the 26°C condition in both strains. Cells of both isolates were also examined by scanning (SEM) and transmission (TEM) electron microscopy (S1 Fig), and cell size was measured at both temperatures (S2 Fig; S2 Table).

**Figure 2.**
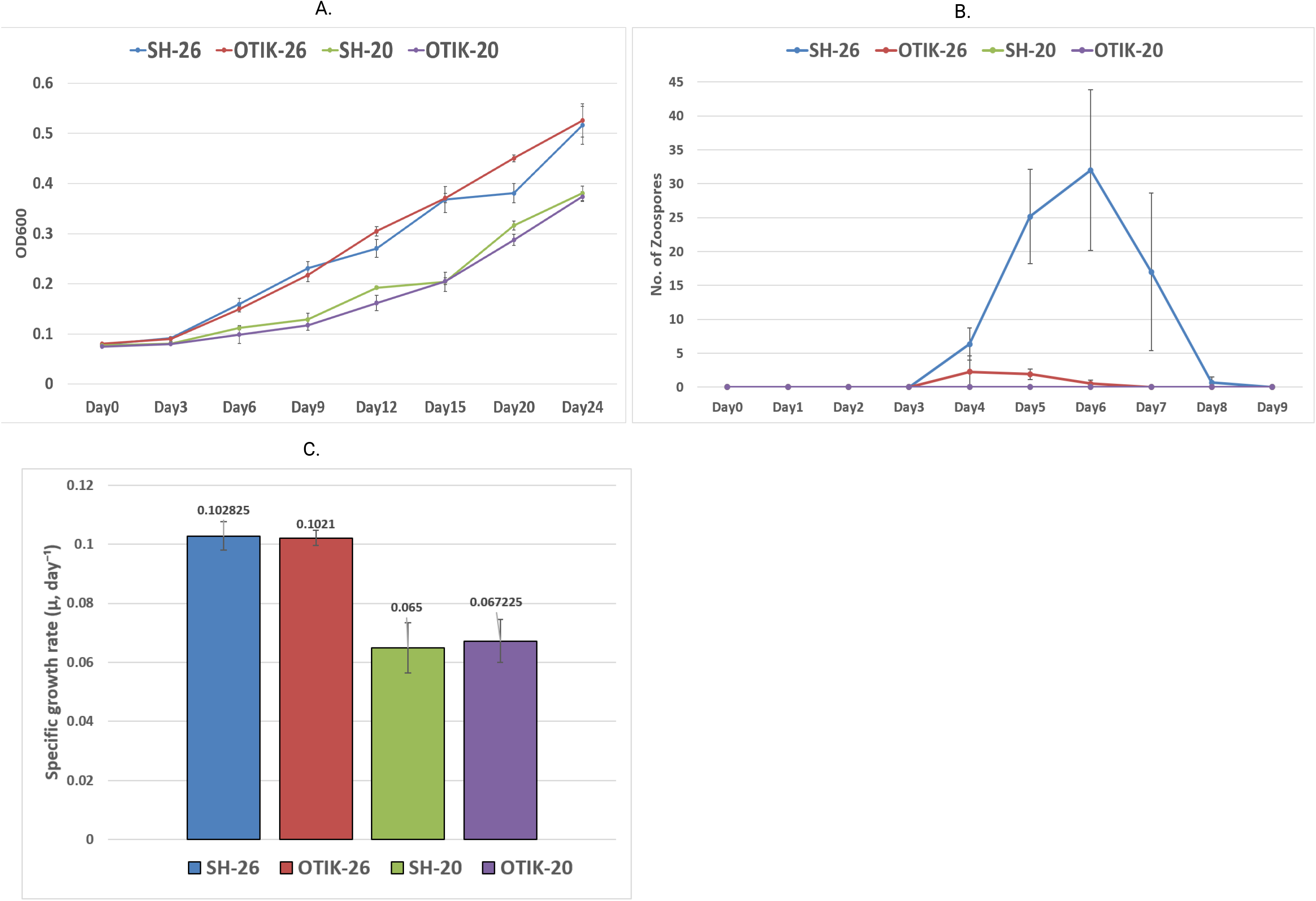
Growth and zoospore production of *Chromera velia* SH and OTI strains at 20°C and 26°C. **(A)** Optical density at 600 nm (OD₆₀₀) of cultures grown in 50 mL f/2 medium in T75 Erlenmeyer flasks under a 12:12 h light:dark photoperiod at 40 μmol photons m⁻² s⁻¹, measured every three days from Day 0 to Day 24. Each line represents the mean of n = 4 biological replicates; error bars represent standard deviation. **(B)** Zoospore counts in the same culture conditions, quantified daily by microscopic counting from Day 0 to Day 9. Each line represents the mean of n = 4 biological replicates; error bars represent standard deviation. **(C)** Specific growth rate (μ) calculated from Day 0 to Day 15 using the formula μ = (ln OD₂ − ln OD₁) / Δt. Bars represent the mean of n = 4 biological replicates; error bars represent standard deviation. A two-way ANOVA (strain × temperature) revealed a significant effect of temperature (F₁,₁₂ = 197.1, p = 8.3 × 10⁻⁹) but not strain (F₁,₁₂ = 0.08, p = 0.78) on specific growth rate, with no significant strain × temperature interaction (F₁,₁₂ = 0.32, p = 0.58).

### Core transcriptome and condition-dependent expression

To establish a baseline of *C. velia* gene expression across strains and temperatures, we identified the genes expressed in each of the four strain-temperature conditions (SH and OTI, each at 20 °C and 26 °C). A gene was considered expressed in a given condition if it exceeded a normalized abundance threshold of CPM ≥ 1 in all three biological replicates for that condition. Across all four conditions, a total of 23,081 genes (the pan-transcriptome) were detectably expressed in at least one condition, of which 15,990 genes (69.3% of the pan-transcriptome) were expressed in all four conditions, forming a stable core transcriptome (Fig 3A, dashed line). The total number of expressed genes per condition varied by strain and temperature. The tropical OTI strain expressed substantially more genes than the temperate SH strain at both temperatures, with 21,198 genes detected in OTI at 20°C and 20,547 at 26°C, compared to 19,018 and 17,259 genes in SH at the same respective temperatures (Fig 3A). The number of expressed genes in OTI varied only modestly between 20°C and 26°C (a decrease of 651 genes, ∼3.1%), whereas SH expressed 1,759 fewer genes at 26°C than at 20°C – a ∼9.2% contraction of the expressed transcriptome (Fig 1A).

**Figure 3.**
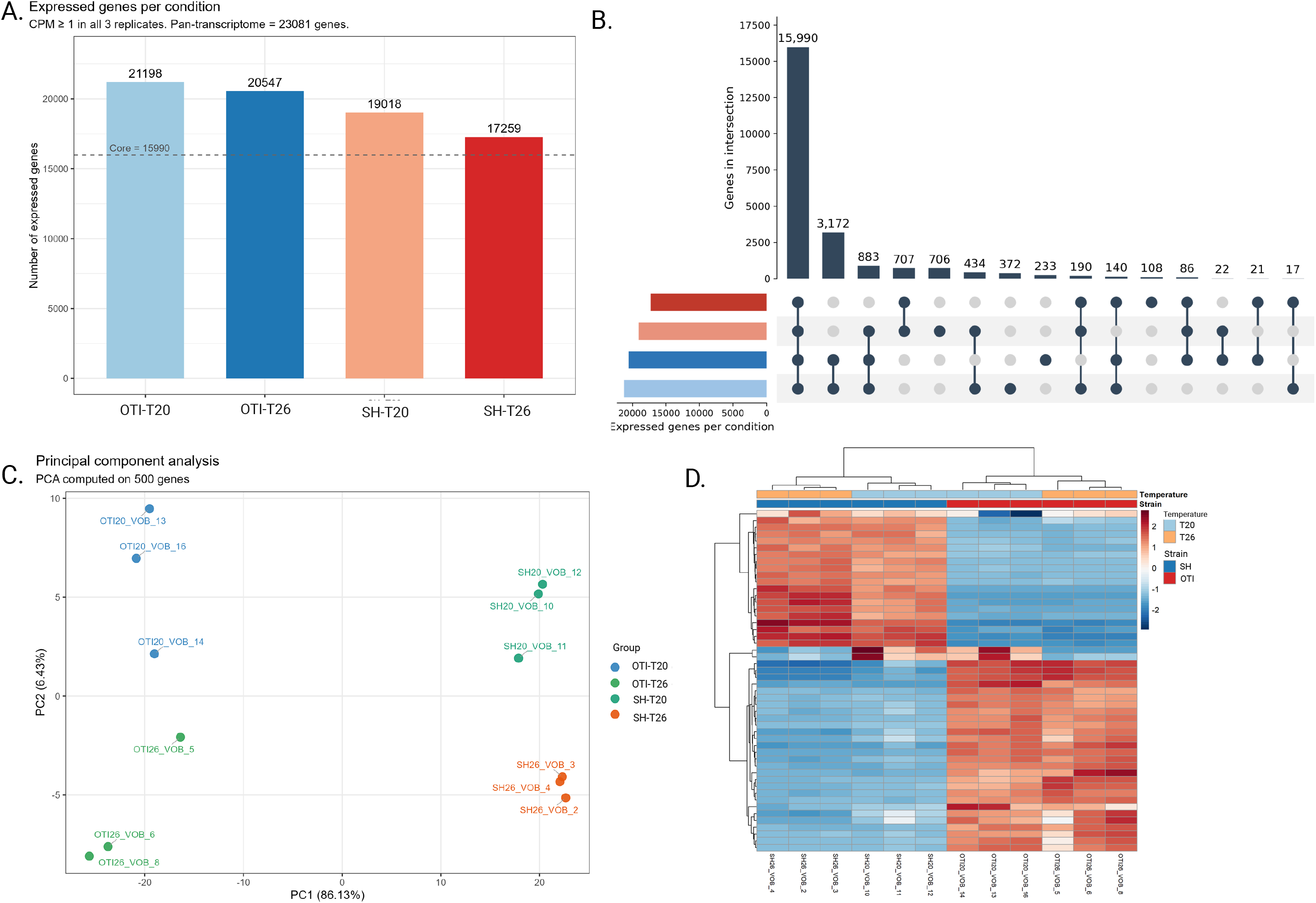
Core transcriptome and condition-specific expression breadth across the four experimental conditions. **(A)** Number of expressed genes per condition, defined as CPM ≥ 1 in all three biological replicates. The dashed line marks the core transcriptome (15,990 genes, expressed in all four conditions); the pan-transcriptome (23,081 genes, expressed in at least one condition) is given in the panel subtitle. **(B)** UpSet plot of intersections between the four condition-specific gene sets. Filled dots indicate which conditions contribute to each intersection; bar heights give the number of genes exclusive to it, and horizontal bars (left) give the total per condition. The largest categories are the 15,990 core genes; 3,172 genes expressed in both OTI conditions but neither SH condition; 883 genes expressed in OTI at both temperatures and SH at 20°C but absent from SH at 26°C; and 707 genes absent only from OTI-T20. **(C)** Principal component analysis of the 12 libraries, computed on variance-stabilising-transformed values of the 500 most variable genes. PC1 (86.13% of variance) separates the two strains; PC2 (6.43%) separates the two temperatures within each strain. **(D)** Hierarchical clustering heatmap of the 50 most variable genes. Values are row-scaled (z-score); red indicates higher and blue lower expression. Annotation bars indicate strain (SH, blue; OTI, red) and temperature (T20, light blue; T26, orange).

The UpSet representation of pairwise and higher-order overlaps among the four condition-specific gene sets (Fig 3B and 3D) further clarifies this pattern. The largest overlap comprises the 15,990 core genes shared by all four conditions. The next largest overlap consists of 3,172 genes expressed in OTI at both temperatures but in neither condition in SH, representing an OTI-specific component of the transcriptome that SH does not express at any temperature. An additional 883 genes are expressed in OTI at both temperatures and in SH at 20°C, but are specifically silent in SH at 26°C. These genes, expressed in SH under cooler conditions, are silenced at the higher temperature, even though OTI maintains their expression under the same thermal regime. Smaller condition-specific and partial-overlap gene sets account for the remaining variation (Fig 3B).

### Overall transcriptome architecture

To resolve the overall transcriptomic relationships among the four experimental conditions, we performed principal component analysis (PCA) on variance-stabilized expression values of the 500 most variable genes (Fig 3C). Strain identity (x-axis) was the dominant axis of variation in the dataset: 86.13% of the total variance was captured by PC1, which clearly separated all OTI samples (left) from all SH samples (right). Temperature (y-axis) contributed a smaller but consistent secondary axis (PC2, 6.43% of the variance), separating samples grown at 20°C from those grown at 26°C within each strain. Biological replicates within each of the four conditions clustered tightly, confirming high reproducibility across the three independent cultures per condition (Fig 3C). Hierarchical clustering of the top 50 genes confirmed the structure observed in the PCA (Fig 3D). The sample dendrogram first grouped samples by strain, then by temperature within each strain, mirroring the PC1–PC2 separation. The heatmap further revealed two clearly demarcated, reciprocally regulated gene blocks: one block was highly expressed in SH but suppressed in OTI, and a second block was highly expressed in OTI but suppressed in SH. Temperature contributed a finer level of substructure within each strain block.

### Strain specific response to similar temperature

To quantify differences in gene transcription between the two strains at each temperature, we performed pairwise DESeq2 contrasts between OTI and SH at 20°C and 26°C separately (padj < 0.05, |log2FC| > 1; SH was set as the reference, so positive log2FC values indicate upregulation in OTI). At 26°C, 8,253 genes were significantly differentially transcribed between the strains: 6,113 (74%) were upregulated in OTI and 2,140 (26%) in SH (Fig 4A). At 20°C, 8,065 genes were significantly differentially transcribed: 5,284 (66%) were upregulated in OTI and 2,781 (34%) in SH (Fig 4B). At both temperatures, OTI upregulated more genes than SH, and this asymmetry was more pronounced at the higher temperature (∼2.9-fold OTI skew at 26°C, compared to ∼1.9-fold at 20°C), indicating that OTI transcripts were upregulated under warmer conditions compared to SH. Overlap between the two strain DEG sets showed that most strain differences appeared across temperatures (Fig 4C). A total of 6,455 genes (65.4% of the union) were differentially transcribed between OTI and SH at both 20°C and 26°C, representing a constitutive component of strain divergence largely independent of growth temperature. The remaining strain DEGs were temperature-specific: 1,798 genes (18.2%) were significant only at 26°C, while 1,610 (16.3%) were significant only at 20°C.

**Figure 4.**
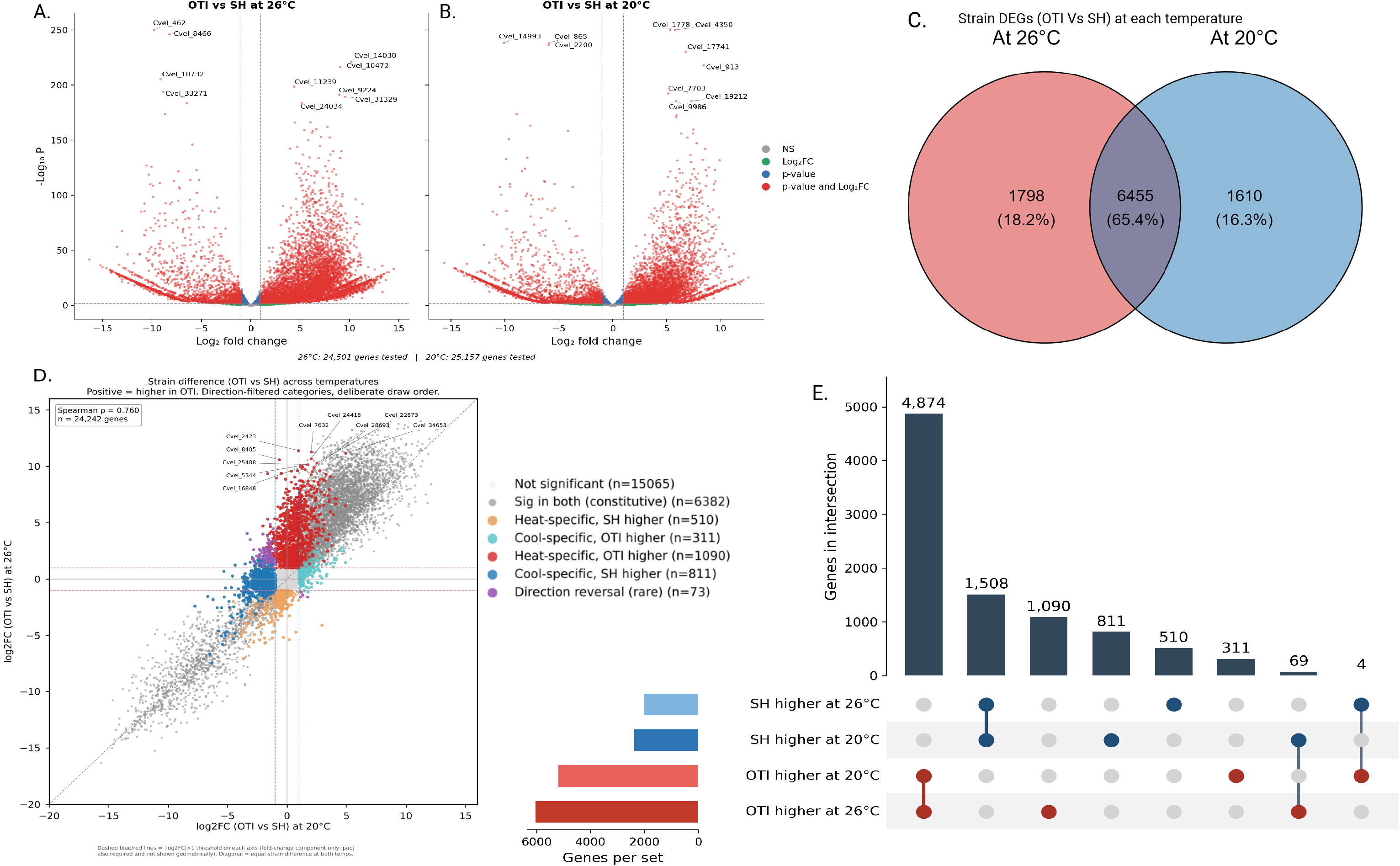
Differential gene expression between the OTI and SH strains at each growth temperature. **(A, B)** Volcano plots of DEGs between OTI and SH at 26°C (A) and 20°C (B), with SH as reference (positive log2FC = higher in OTI). Dashed lines mark the effect-size (|log2FC| > 1) and significance (padj < 0.05) thresholds; genes meeting both are red, significance only blue, effect-size only green, neither grey. 8,253 of 24,524 genes tested were significant at 26°C (6,113 higher in OTI; 2,140 higher in SH), and 8,065 of 25,187 at 20°C (5,284 higher in OTI; 2,781 higher in SH). **(C)** Venn diagram of strain DEGs at the two temperatures: 6,455 genes (65.4%) were significant at both, representing the constitutive component of strain divergence, while 1,798 (18.2%) and 1,610 (16.3%) were significant only at 26°C or only at 20°C. **(D)** Strain difference (OTI vs SH log2FC) at 20°C versus 26°C for all 24,242 genes with a log2FC at both temperatures (Spearman ρ = 0.760). The dotted diagonal marks an equal strain difference at both temperatures. Genes are colored by significance category (padj < 0.05, |log2FC| > 1, direction-matched at each temperature). The bars show the number of genes higher in each strain at each temperature **(E)** UpSet plot partitioning all 9,177 genes significant at one or both temperatures. Bar heights give the number of genes exclusive to each intersection. The four leftmost bars comprise the 6,455 constitutive genes from (C): 4,874 higher in OTI at both temperatures, 1,508 higher in SH at both, and rare direction reversals (69 and 4). The remainder are the temperature-specific sets described in (D). **Differential expression was tested with DESeq2 (padj < 0.05, |log2FC| > 1); n = 3 biological replicates per condition.**

To determine whether the strain difference for each gene remained stable in magnitude or was amplified at one temperature, we plotted the log2FC values of all expressed genes at 20°C against their corresponding values at 26°C (Fig 4D). Most genes clustered closely along the unity diagonal, indicating that the strain difference for the majority of genes was maintained in both direction and magnitude across the two temperatures, consistent with the constitutive component identified above. However, a distinct off-diagonal cluster of 1,090 genes was shifted upward from the diagonal (red, Y > 0): these genes showed little or no strain transcriptional variance at 20°C but were strongly upregulated in OTI relative to SH at 26°C, representing a heat-specific OTI gene set. A reciprocal but smaller cool-specific SH cluster of 811 genes was also observed (blue, X < 0). Strain transcriptomic differences that reversed direction between the two temperatures were rare: only 69 genes were transcribed higher in SH at 20°C and higher in OTI at 26°C, and only 4 genes showed the opposite pattern, confirming that strain transcriptomic differences rarely reverse direction across the tested thermal range. Together, these analyses demonstrate that strain transcriptomic divergence in *Chromera* is large, predominantly constitutive, and asymmetrically expanded at the higher temperature in the OTI strain but not in SH.

### Temperature-specific gene transcription in SH and OTI

To characterize how each strain transcriptionally responded to a change in temperature, we performed within-strain DESeq2 contrasts comparing 26°C to 20°C (with T20 set as the reference, so positive log2FC values indicate higher expression at 26°C). In SH, 4,511 genes responded significantly to the temperature shift, of which only 692 (15.3%) were upregulated at 26°C, while 3,819 (84.7%) were downregulated (Fig 5A). In OTI, 4,089 genes responded significantly, with 1,647 (40.3%) upregulated and 2,442 (59.7%) downregulated (Fig 5B). The two strains thus differed substantially in the nature of their transcriptional response to temperature: SH’s response was strongly suppression-dominated, with downregulated genes outnumbering upregulated genes by more than fivefold, whereas OTI’s transcriptional response was considerably more balanced. The magnitude of the response also differed markedly between strains; SH thermal DEGs reached much higher significance values and larger effect sizes (-log10 padj up to ∼150) than OTI DEGs (-log10 padj up to ∼30), indicating a more extreme inhibition of transcription in SH upon exposure to the higher temperature. This pattern of broad, large-magnitude suppression is the canonical transcriptomic signature of acute thermal stress, consistent with 26°C lying above SH’s preferred thermal range, whereas OTI’s more measured and balanced response is consistent with acclimation within its tolerated range. Overlap between the two strains’ thermal-response gene sets showed that 1,614 genes (23.1% of the union) responded to temperature in both strains, while 2,897 genes (41.5%) responded only in SH and 2,475 genes (35.4%) only in OTI (Fig 5C). The shared set of 1,614 genes represents a conserved core thermal response common to both strains.

**Figure 5.**
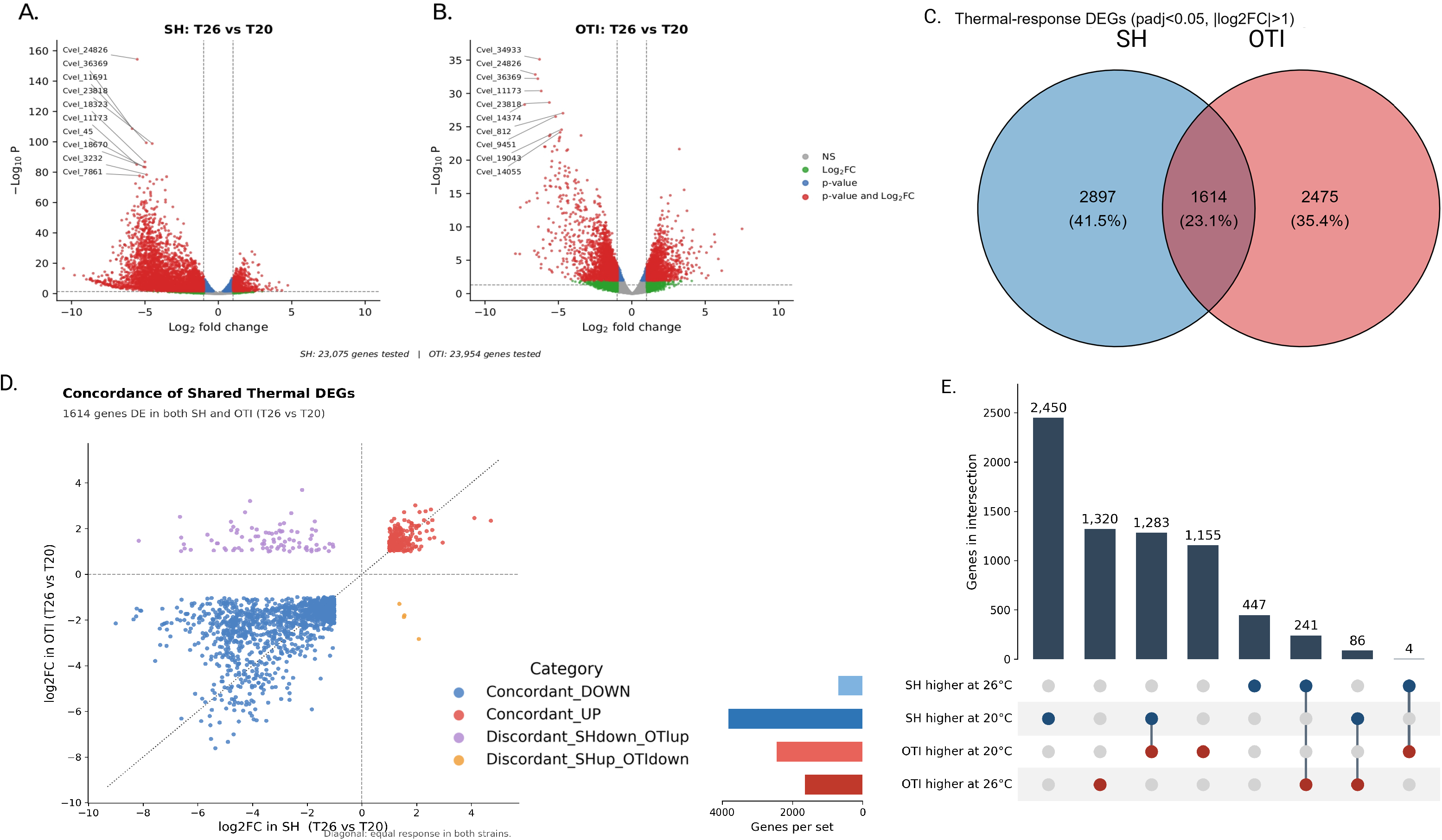
Within-strain thermal-response transcriptomes differ markedly in magnitude and balance between SH and OTI. **(A, B)** Volcano plots of DEGs between 26°C and 20°C within SH (A) and OTI (B), with 20°C as reference (positive log2FC = higher at 26°C). Thresholds and colour coding as in Figure 4A. In SH, 4,511 genes were significantly differentially expressed (692 up, 3,819 down at 26°C), the response being strongly dominated by suppression. In OTI, 4,089 genes were significant (1,647 up, 2,442 down at 26°C), giving a more balanced response. **(C)** Venn diagram of significant thermal-response genes in the two strains. 1,614 genes (23.1%) responded to temperature in both strains, representing a conserved core thermal response; 2,897 (41.5%) responded only in SH and 2,475 (35.4%) only in OTI. **(D)** Thermal-response log2FC (26°C vs 20°C) in SH (x-axis) versus OTI (y-axis), restricted to the 1,614 genes responding significantly in both strains (C). Dashed grey lines mark log2FC = 0; the dotted diagonal marks an identical response in both strains. Genes are coloured by concordance: 1,283 (79.5%) concordantly downregulated at 26°C in both strains (blue); 241 (14.9%) concordantly upregulated (red); 86 (5.3%) discordant, being downregulated at 26°C in SH but upregulated in OTI (purple) — that is, each strain expressed these genes more highly at its own native temperature; and 4 (0.2%) showing the reciprocal discordant pattern (orange). The 20 discordant genes diverging most from the diagonal are labelled.**(E)** UpSet plot partitioning all 6,986 thermal-response genes by strain and direction. Bar heights give the number of genes exclusive to each intersection. The four single-set bars are the strain-unique responses (2,450 and 447 in SH; 1,155 and 1,320 in OTI), and the four paired bars are the shared responses from (D): 1,283 concordantly higher at 20°C, 241 concordantly higher at 26°C, and the discordant sets of 86 and 4 genes. **Differential expression was tested with DESeq2 (padj < 0.05, |log2FC| > 1); n = 3 biological replicates per condition.**

To determine whether transcription of these shared thermal-response genes was regulated in the same direction in both strains, we plotted each gene’s log2FC in SH against its log2FC in OTI (Fig 5D). Most shared genes responded concordantly: 1,283 genes (79.5%) were downregulated at 26°C in both strains, and 241 genes (14.9%) were upregulated at 26°C in both strains, together accounting for 94.4% of the shared set. However, 86 genes (5.3%) showed discordant regulation, being downregulated at 26°C in SH but upregulated at 26°C in OTI – that is, each strain expressed these genes more highly at its own native temperature. The opposite discordant pattern (upregulated in SH but downregulated in OTI) was observed for only 4 genes (0.2%). This approximately 20-fold asymmetry between the two discordant categories indicates that the discordance is unlikely to reflect random noise, but instead represents directional regulatory divergence between the strains.

### Gene ontology enrichment of strain- and thermal-response gene sets

To resolve the functional categories underlying the differentially expressed genes, we performed GO enrichment separately on gene sets derived from two independent contrasts. For the strain comparison, we took the genes differentially expressed between OTI and SH at each temperature and partitioned them by direction, yielding the genes more highly expressed in OTI and those more highly expressed in SH at 20°C (n = 311 and n = 811 significant DEGs, respectively) and at 26°C (n = 1,090 and n = 510) (Fig 4E). For the thermal response, we took each strain’s within-strain contrast of 26°C against 20°C and partitioned it by direction, yielding the genes upregulated at each temperature in OTI (n = 1,320 up at 26°C; n = 1,155 up at 20°C) and in SH (n = 447 up at 26°C; n = 2,450 up at 20°C) (Fig 4E). Enrichment was assessed against a Benjamini–Hochberg FDR threshold of 0.05, and terms not passing this threshold are shown in grey (Fig 6).

**Figure 6.**
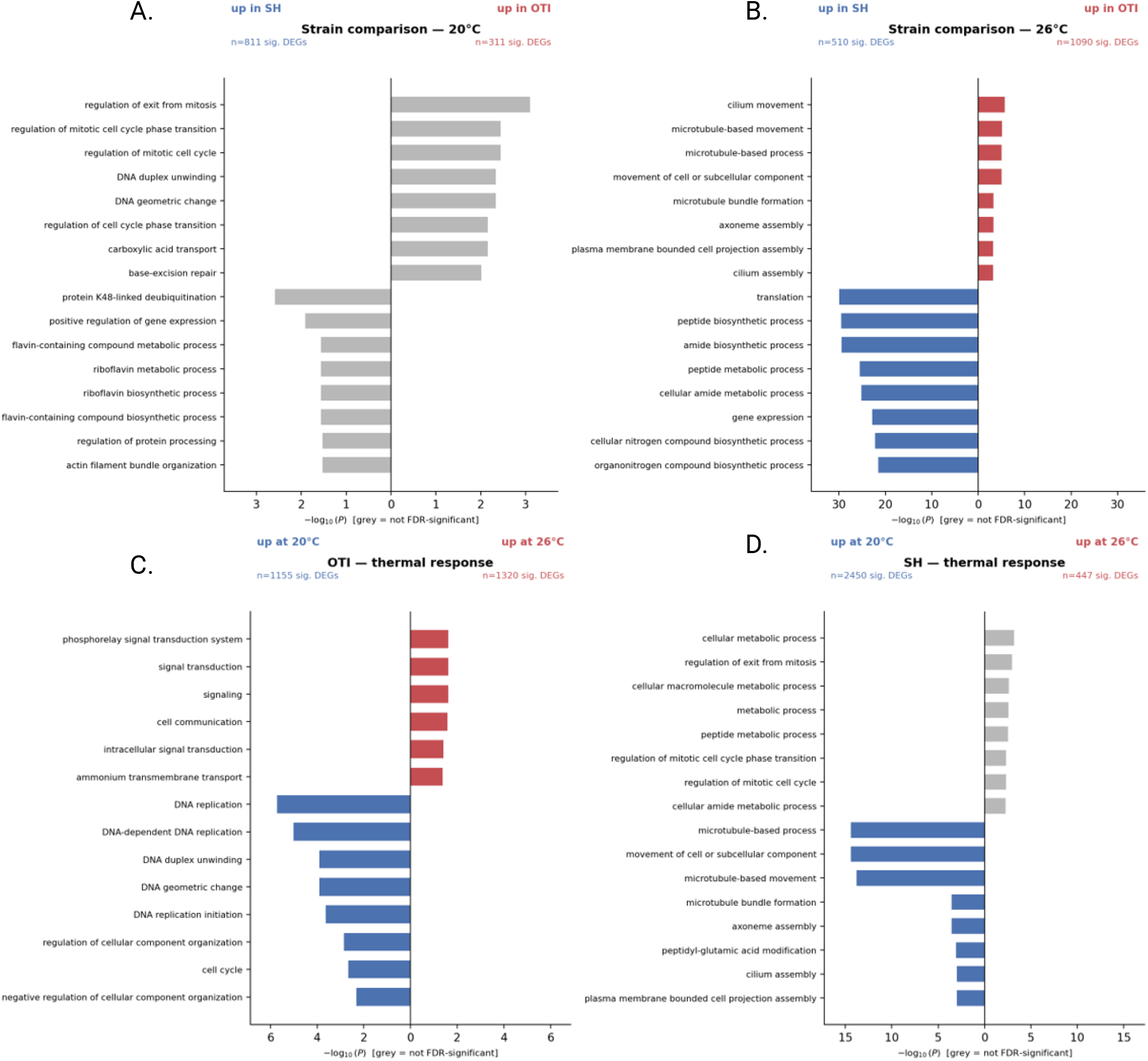
Gene ontology enrichment of strain- and thermal-response gene sets. **(A, B)** GO terms enriched in strain-comparison gene sets at 20°C (A) and 26°C (B), plotted as diverging bars (up in SH, blue leftward; up in OTI, red rightward). Bar length gives −log10(P); coloured bars indicate FDR-significant terms (BH-adjusted P < 0.05), grey bars indicate the most nominally significant terms in gene sets that returned no FDR-significant enrichment. The number of significant DEGs contributing to each direction is shown above each panel. At 20°C, neither direction returned FDR-significant terms despite the underlying DEGs. At 26°C, OTI-elevated genes were enriched for cilium and microtubule terms, and SH-elevated genes for translation and biosynthesis terms. **(C, D)** GO terms enriched in the within-strain thermal-response gene sets for OTI (C) and SH (D), plotted as up at 20°C (blue, leftward) versus up at 26°C (red, rightward). Colour coding as in (A, B). In OTI, genes upregulated at 26°C were enriched for signal transduction, and genes upregulated at 20°C for DNA replication and cell cycle. In SH, genes upregulated at 20°C were enriched for microtubule and cilium terms, while genes upregulated at 26°C returned no FDR-significant enrichment. For each panel, the top ranked GO terms (BH-adjusted P < 0.05, or top nominal-P where no term reached FDR significance) are shown; DEG counts and total tested gene numbers are as reported in Figures 4 and 5.

In the strain comparison at 20°C, no GO term reached FDR significance in either direction despite the underlying DEGs, and all displayed terms are grey (Fig 6A). At 26°C, both directions returned significant enrichment (Fig 6B): genes more highly expressed in OTI were enriched for cilium- and microtubule-based terms (cilium movement, microtubule-based movement and process, axoneme assembly, cilium assembly), while genes more highly expressed in SH were enriched for translation and biosynthesis terms (translation, peptide and amide biosynthetic process, gene expression, nitrogen compound biosynthetic process), the latter reaching −log10(P) values above 30. In the thermal-response panels, OTI’s genes upregulated at 26°C were enriched for signal transduction terms (phosphorelay signal transduction, signaling, cell communication, ammonium transmembrane transport), and its genes upregulated at 20°C were enriched for DNA replication and cell-cycle terms (DNA replication, DNA duplex unwinding, cell cycle) (Fig 6C). For SH, genes upregulated at 20°C were enriched for microtubule- and cilium-based terms (microtubule-based movement and process, axoneme assembly, cilium assembly), while genes upregulated at 26°C returned no FDR-significant terms and are shown in grey (Fig 6D).

### Gene-level expression across pathways

To examine expression at the level of individual genes rather than enriched categories, we annotated the DEGs and assembled four functional gene sets — photosynthesis/chlorophyll, ribosome/translation, lipid-related, and cilium/microtubule-motility — displaying each as a radial heatmap in which four concentric rings show, from inner to outer, the strain difference (OTI vs SH) at 20°C and at 26°C and the within-strain thermal response (26°C vs 20°C) for SH and for OTI. Cells are colored by log2 fold-change where significant (FDR < 0.05), grey where the comparison was not significant, and black where the gene was not detected (Fig 7). For the photosynthesis/chlorophyll set (18 genes; Fig 7A), the strain rings were mixed in direction: at 20°C, 9 of 18 genes were significant (4 higher in OTI, 5 higher in SH), and at 26°C, 12 of 18 were significant (3 higher in OTI, 9 higher in SH). The SH thermal ring showed the most significant cells of the four rings (14 of 18 genes), the majority upregulated at 26°C (10 genes), while the OTI thermal ring showed 8 of 18 genes significant, split evenly between up- and downregulation. For the ribosome/translation set (18 genes; Fig 7B), the two strain rings (inner two) were dominated by negative values: 15 of 18 genes were significant at 20°C and 17 of 18 at 26°C, and in both rings the large majority were lower in OTI than in SH (13 and 15 genes negative, respectively), with two genes positive in each. The two thermal rings (outer two) showed comparatively little signal, with 5 of 18 genes significant in each of the SH and OTI thermal responses, predominantly downregulated at 26°C. For the lipid-related set (18 genes; Fig 7C), the two strain rings were dominated by positive values: 15 of 18 genes were significant at 20°C and 16 of 18 at 26°C, with most genes higher in OTI than in SH (12 and 14 genes positive, respectively). The SH thermal ring was the inverse, with 16 of 18 genes significant and nearly all downregulated at 26°C (15 genes), whereas the OTI thermal ring showed only 7 of 18 genes significant, mixed in direction. For the cilium/microtubule-motility set (18 genes; Fig 7D), the strain rings differed sharply between temperatures: at 20°C only 1 of 18 genes was significant, whereas at 26°C all 18 genes were significant and every one was higher in OTI than in SH. The thermal rings were also asymmetric between strains: 16 of 18 genes were significant in the SH thermal response and all were downregulated at 26°C, while only 1 of 18 genes was significant in the OTI thermal response.

**Figure 7.**
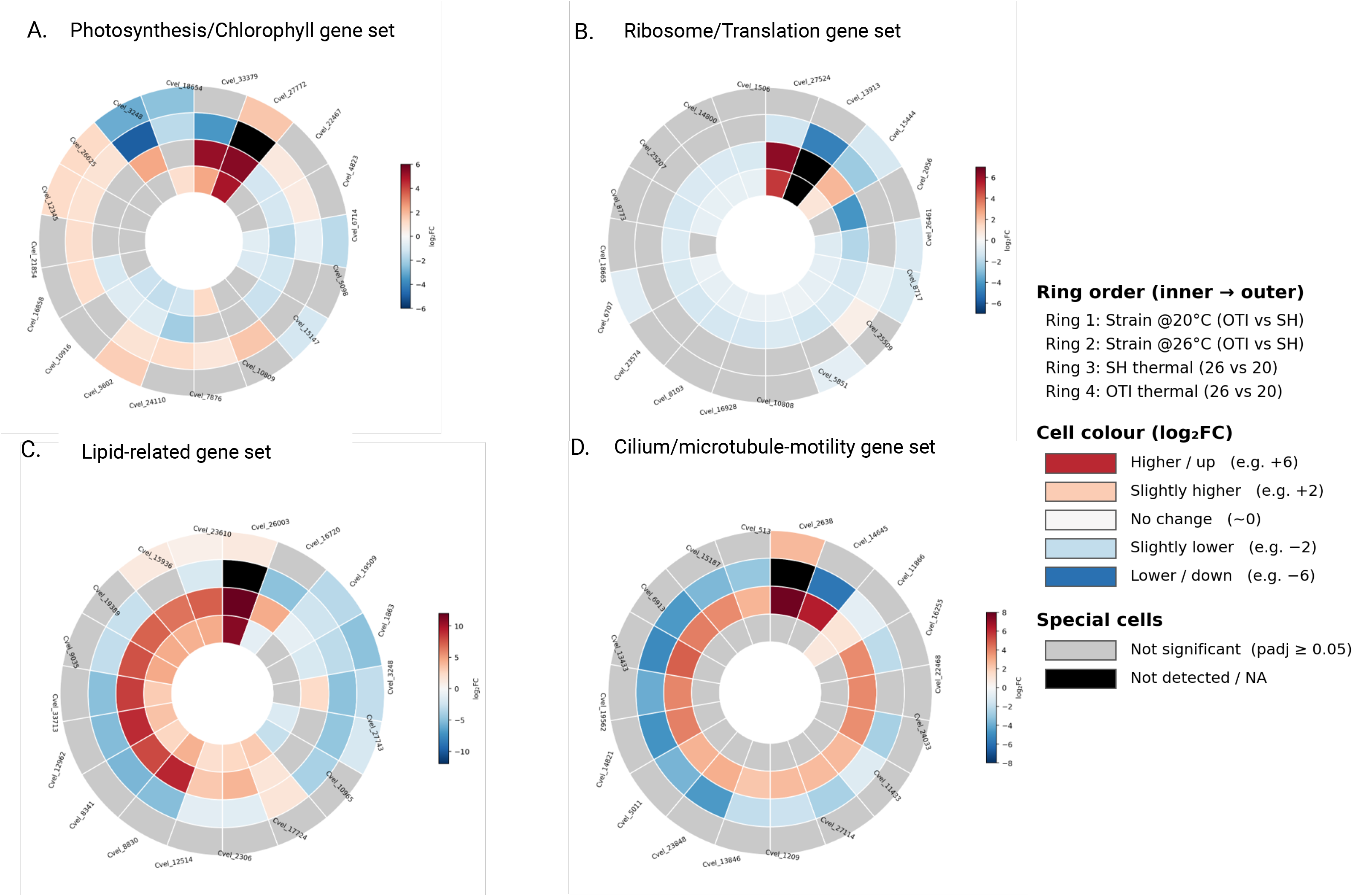
Gene-level expression patterns across four functional gene sets. Radial heatmaps of the top significant genes belonging to four functional categories: (A) photosynthesis / chlorophyll, (B) ribosome / translation, (C) lipid-related, and (D) cilium / microtubule-motility. Within each panel, the four concentric rings show, from inner to outer, the strain difference (OTI vs SH; positive = higher in OTI) at 20°C and 26°C, followed by the within-strain thermal response (26°C vs 20°C; positive = higher at 26°C) for SH and for OTI. Cells are coloured by log2 fold-change where the underlying comparison was significant (padj < 0.05, DESeq2); grey cells indicate non-significant comparisons and black cells indicate genes not detected. Colour scales are independent per panel. Genes are ordered by hierarchical clustering on their combined pattern across all four rings, and Chromera velia (Cvel) gene IDs are shown at the perimeter. Gene selection: top 18 genes per category ranked by combined significance across comparisons, drawn from the pool of annotated genes matching each functional keyword set. Ring assignment and colour coding are shown in the shared key (right).

### Fatty acid composition of membrane and storage lipid classes

To resolve the lipidomic response at the level of individual acyl chains, we determined the fatty acid composition of each lipid class separately in both isolates at both temperatures. Classes were assigned to a membrane pool (MGDG, DGDG, SQDG+PG, DGTS, DGTA) and a storage pool (TG, DG). Because the analysis was performed on a fixed mass of lipid extract, the total fatty acid signal recovered per replicate was constant across all four conditions (916,700 ± 10,600 to 952,000 ± 2,000), and all values are therefore proportions of the extracted lipid rather than quantities per cell or per unit biomass. Ionisation response differs substantially between lipid classes in this method — DGTS accounted for 96.8–98.0 % of the membrane pool signal, with MGDG at 0.15–1.03 % — so comparisons are made within a class across conditions and not between classes. Total lipid content, measured gravimetrically at 20 °C, was higher in SH than in OTI (13.05 ± 1.94 % versus 10.42 ± 1.86 % of dry mass, p = 0.042).

### Lipid class composition is indistinguishable between the isolates at 20 °C but diverges at 26 °C

Principal component analysis of centered log-ratio transformed class composition separated the four conditions along PC1 (72.7 % of variance) and PC2 (19.3 %), with the two isolates overlapping completely at 20 °C and forming distinct, non- overlapping clusters at 26 °C (Fig 8A). This pattern was quantified by the ratio of membrane to storage fatty acids (Fig 8B). At 20°C the two isolates were effectively identical (OTI 0.158 ± 0.066; SH 0.163 ± 0.025; p = 0.88). At 26 °C they had moved in opposite directions, OTI to 0.220 ± 0.011 and SH to 0.078 ± 0.050 (p = 0.034), yielding a significant strain × temperature interaction (p = 0.013). The same interaction was present in the membrane pool considered alone (p = 0.008) and was marginal in the storage pool (p = 0.054), indicating that the divergence was driven primarily by the membrane side of the allocation. Neither within-strain change reached significance on its own (SH −52 %, p = 0.080; OTI +39 %, p = 0.157), so the result is best expressed as a difference in how the two isolates responded rather than as two independent responses.

**Fig 8.**
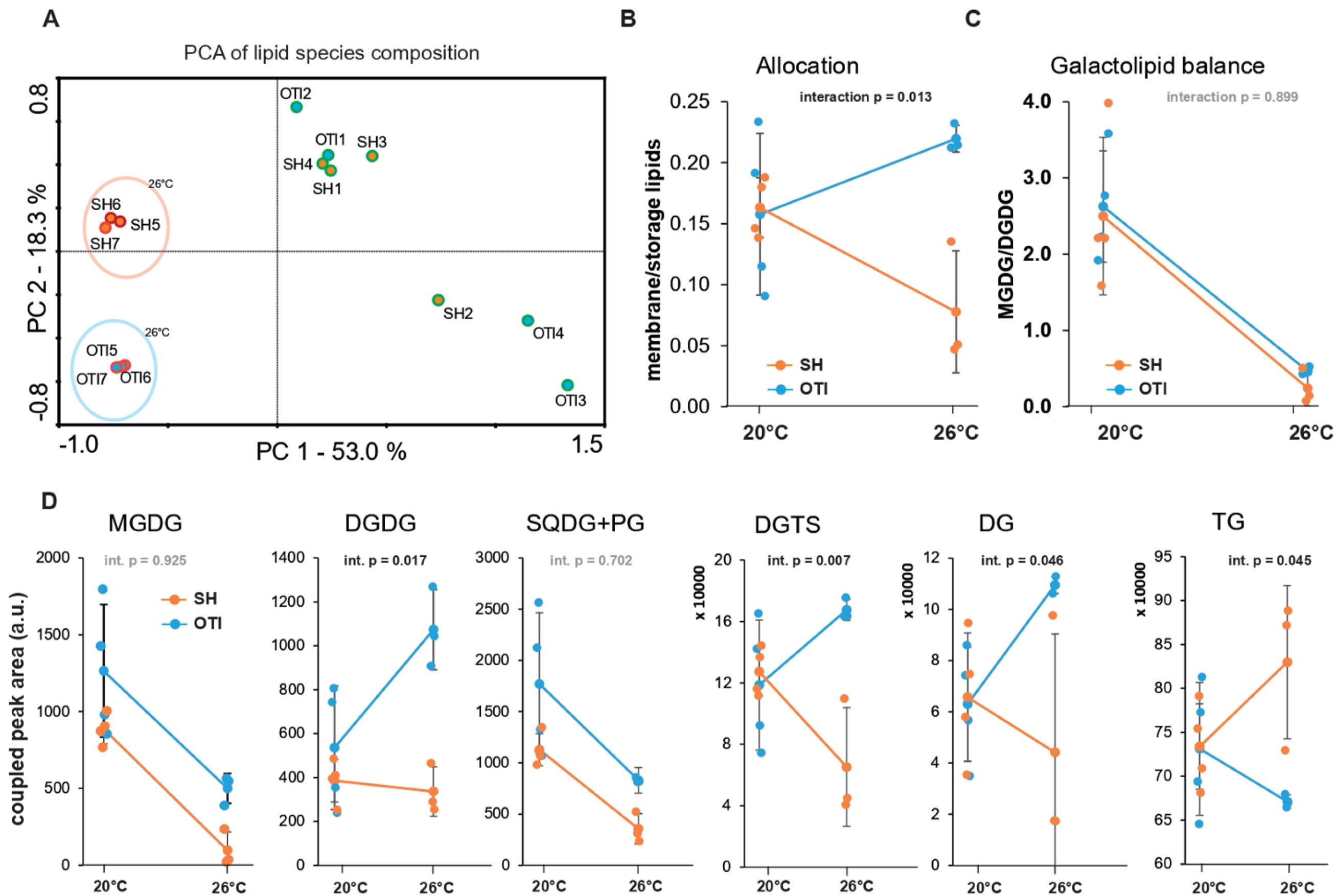
Lipid class composition is indistinguishable between the two isolates at 20 °C and diverges at 26 °C. **(A)** Principal component analysis the 204 intact lipid species detected by LC-MS/MS; PC1 and PC2 account for 53.0 % and 18.3 % of the variance. Samples are labelled by isolate and replicate: OTI1–OTI4 and SH1–SH4 were grown at 20 °C, OTI5–OTI7 and SH5–SH7 at 26 °C. Point color denotes growth temperature. Shaded ellipses mark the 26°C replicates of each isolate and are drawn for visual grouping only; they are not confidence regions. The 20 °C samples of the two isolates intermix, whereas the 26 °C samples from two separate, tight clusters. **(B)** Ratio of membrane to storage fatty acids. Membrane classes are MGDG, DGDG, SQDG+PG, DGTS and DGTA; storage classes are TG and DG. The isolates are indistinguishable at 20 °C (0.158 ± 0.066 in OTI, 0.163 ± 0.025 in SH; p = 0.88) and differ at 26 °C (0.220 ± 0.011 and 0.078 ± 0.050; p = 0.034). **(C)** MGDG:DGDG ratio. The ratio falls in both isolates, from 2.63 ± 0.73 to 0.47 ± 0.05 in OTI (p = 0.009) and from 2.50 ± 1.03 to 0.24 ± 0.23 in SH (p = 0.019), with no difference between the isolates at either temperature. This component of the thermal response is shared. **(D)** Abundance of the six individual lipid classes, as coupled peak area. Note the differing y-axis scales and multipliers between panels; the TG axis does not begin at zero. MGDG declines in both isolates (SH −89 %, p < 0.001; OTI −60 %, p = 0.034), as does SQDG+PG (SH −68 %, p = 0.002; OTI −53 %, p = 0.073). DGDG doubles in OTI (p = 0.028) while remaining unchanged in SH (p = 0.58). DG rises in OTI (p = 0.023). TG shows no significant within-strain change in SH (+13 %, p = 0.19). *Lines connect group means, error bars show SD and points are individual replicates (n = 4 at 20 °C, n = 3 at 26 °C). Strain × temperature interaction p-values are from two-way ANOVA (type II sums of squares); pairwise comparisons in the text use two-sided Welch’s t-tests, without correction for multiple testing. Values are proportions of a fixed mass of lipid extract and are compositional, so they can be compared within a class across conditions but not between classes.

At the level of individual classes (Fig 8D), MGDG declined in both isolates (SH −89 %, p < 0.001; OTI −60 %, p = 0.034) with no interaction (p = 0.92), as did SQDG+PG (SH −68 %, p = 0.002; OTI −53 %, p = 0.073; interaction p = 0.70). DGDG, by contrast, doubled in OTI (p = 0.028) while remaining unchanged in SH (p = 0.58; interaction p = 0.017), and DGTS, the dominant membrane class in this dataset, showed a significant interaction (p = 0.007) with the two isolates again moving in opposite directions. Among the storage classes, DG rose in OTI (+74 %, p = 0.023) and TG showed a significant interaction (p = 0.045) but no significant within-strain change in SH (+13 %, p = 0.19). The shift in allocation in SH therefore reflects a contraction of the membrane pool rather than accumulation of triacylglycerol.

### Lipid profile is a conserved component of the thermal response

Two features of the response were shared by both isolates. The MGDG:DGDG ratio fell from 2.63 ± 0.73 to 0.47 ± 0.05 in OTI (p = 0.009) and from 2.50 ± 1.03 to 0.24 ± 0.23 in SH (p = 0.019), with no interaction (p = 0.90) and no difference between the isolates at either temperature (Fig 8C). Both isolates also withdrew eicosapentaenoic acid (20:5n-3) from their membrane lipids and recovered it in their storage lipids (Figs 9A and 9D): membrane 20:5n-3 fell by 93 % in SH (p < 0.001) and 62 % in OTI (p = 0.050), while storage 20:5n-3 rose by 92 % (p = 0.002) and 139 % (p = 0.002), respectively. The interaction was significant on the membrane side (p = 0.012) but not on the storage side (p = 0.195), indicating that the destination of the withdrawn chains was common to both isolates even though the extent of withdrawal differed. Arachidonic acid (20:4n-6) behaved similarly, rising in the storage pool in both isolates (SH +44 %, p = 0.003; OTI +42 %, p = 0.048) with no interaction (p = 0.95).

**Fig 9.**
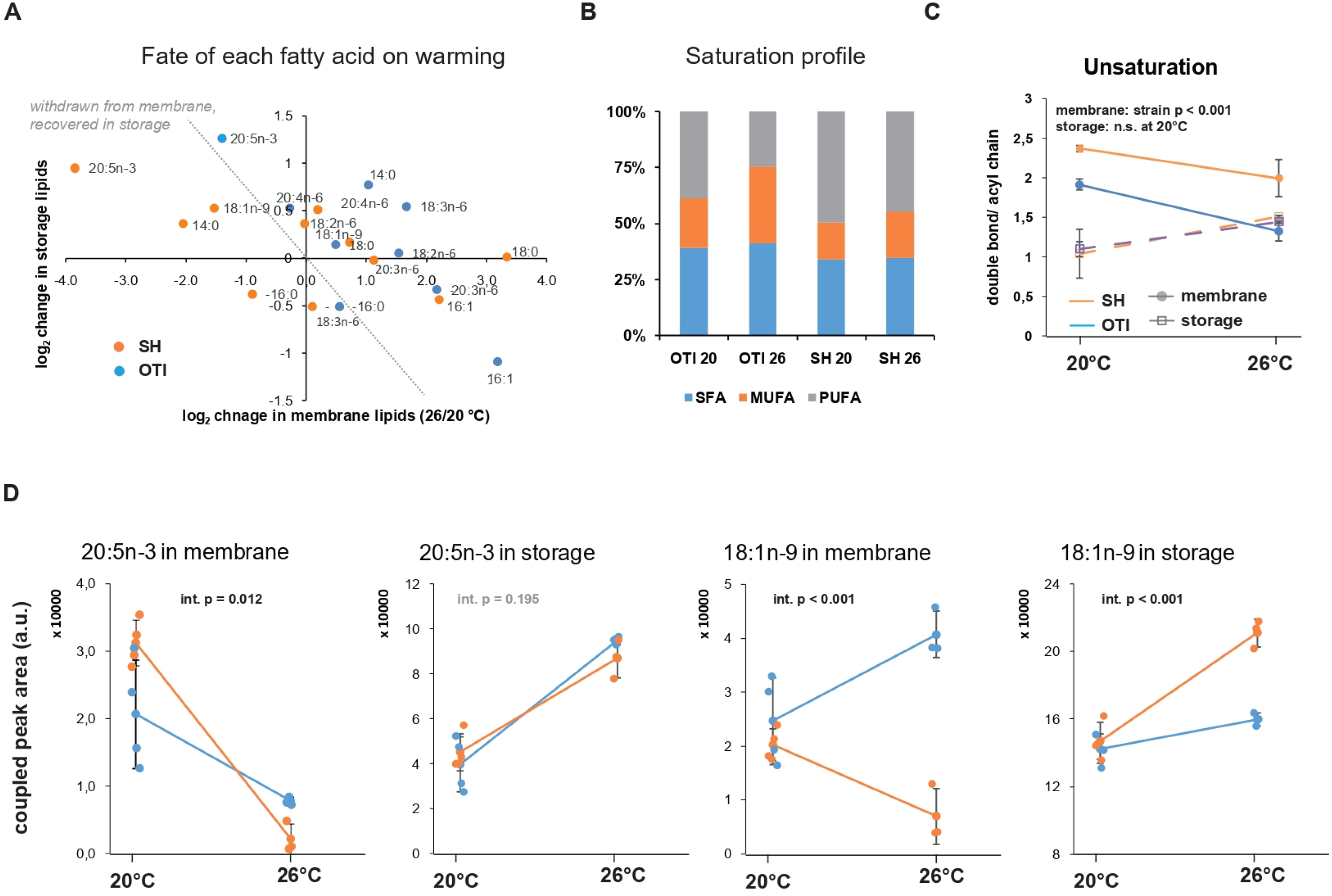
Both isolates withdraw polyunsaturated chains from their membranes at 26 °C, but only OTI replaces them. **(A)** Fate of each fatty acid on warming: log2 change (26 °C/20 °C) in the storage pool against the log2 change in the membrane pool, one point per fatty acid and isolate. Points near the anti-diagonal fell in one pool and rose in the other, consistent with redistribution between pools; 20:5n-3 does so in both isolates. The panel is a descriptive plot of group means with no statistical test. **(B)** Membrane fatty acid saturation profile, as proportions of saturated (SFA), monounsaturated (MUFA) and polyunsaturated (PUFA) chains. **(C)** Mean number of double bonds per acyl chain in the membrane (solid lines) and storage (dashed lines) pools. **(D)** 20:5n-3 and 18:1n-9 in the membrane and storage pools, as coupled peak area (a.u.). Panels B and C are calculated within each pool, whereas D is not normalised to pool size. Lines connect group means. Error bars show SD (C, D) and points in (D) are individual replicates; n = 4 biological replicates at 20 °C and n = 3 at 26 °C. Strain × temperature interaction p-values are from two-way ANOVA (type II sums of squares); pairwise comparisons in the text use two-sided Welch’s t-tests, without correction for multiple testing. Values are proportions of a fixed mass of lipid extract and are therefore compositional.

### OTI membranes are constitutively less unsaturated

The two isolates differed in membrane acyl composition at both temperatures (Figs 9B and 9C). At 20 °C the membrane pool of OTI carried 1.92 ± 0.07 double bonds per acyl chain, compared with 2.37 ± 0.04 in SH (p = 0.0001), and polyunsaturated chains accounted for 38.9 % of OTI membrane fatty acids versus 49.5 % in SH (p = 0.0002). Both isolates became less unsaturated at 26 °C (OTI 1.32 ± 0.12; SH 2.00 ± 0.24; effect of temperature p < 0.001), and the separation between them persisted (p = 0.023) with no strain × temperature interaction (p = 0.14). The difference was specific to the membrane pool: storage lipids did not differ between isolates at 20°C (1.04 ± 0.31 versus 1.10 ± 0.09; p = 0.73). OTI therefore maintains a constitutively less unsaturated membrane, and this appears to be an inherent property of the isolate rather than a response to the temperature imposed here.

### The isolates differ in the fate of newly synthesized monounsaturated chains

Plotting the thermal change of each fatty acid in the membrane pool against its change in the storage pool (Fig 9A) separates the shared from the divergent components of the response. Chains withdrawn from the membrane and recovered in storage, of which 20:5n-3 is the clearest example, fall near the anti-diagonal in both isolates. Oleic acid (18:1n-9) does not. Both isolates held more 18:1n-9 at 26 °C than at 20 °C (SH +30 %, p = 0.002; OTI +20 %, p = 0.021), but its distribution differed sharply (Fig 9D). In OTI, membrane 18:1n-9 rose by 65 % (p = 0.022) and storage 18:1n-9 by only 12 % (p = 0.020). In SH, membrane 18:1n-9 fell by 65 % (p = 0.029) while storage 18:1n-9 rose by 44 % (p < 0.001). The strain × temperature interaction was significant on both pools (membrane p < 0.001; storage p < 0.001), and the two isolates, indistinguishable in membrane 18:1n-9 at 20 °C (p = 0.36), differed by sixfold at 26 °C (p = 0.001). Palmitic acid (16:0) followed the same pattern in the membrane pool, with a significant interaction (p = 0.007) and a significant difference between the isolates at 26 °C (p = 0.018) but not at 20 °C (p = 0.66).

Taken together, the lipidomic data describe one response common to both isolates: withdrawal of polyunsaturated chains from the membrane into storage lipid, with a coordinated shift in the galactolipid balance and one response that distinguishes them: OTI directs monounsaturated chains into its membrane lipids and expands its membrane pool, whereas SH directs them into storage and its membrane pool contracts.

## Discussion

Coral reefs support marine biodiversity and provide coastal protection, fisheries, and livelihoods for millions of people [35]. Reef-building corals depend on a symbiosis with photosynthetic algae that live inside their tissues and supply most of the energy they need [36]. The algae associated with corals span a spectrum of relationships, from essential mutualists to opportunists. *Chromera velia*, first isolated from reef corals, sits at the uncertain end of this spectrum: rather than supporting its host, it can infect coral larvae and behave as a parasite [14,15], even though it was first isolated from reef corals using the same methods routinely used to isolate symbiotic Symbiodiniaceae [10]. Beyond its association with corals, *C. velia* has a far broader ecological range than its original isolation suggested. Environmental sequencing studies have detected it worldwide and not necessarily in association with corals [17,18], implying that the species persists across a wide range of habitats and temperature conditions. This is corroborated by 18S rDNA phylogeny of *C. velia* isolate from one tree island (OTI) alongside environmental amplicon sequence variants (ASV’s) from geographically diverse sites (Bahamas, Canada, Australia), which grouped within the same broader clade (S3 Fig) [18]. This raises a fundamental question about how a single species copes with such environmental variation: are geographically separated populations physiologically equivalent, or have they diverged in ways that reflect their local conditions? The ease of cultivation[10,11] and the availability of nuclear and organellar genome sequences [37–39]make *C. velia* a powerful experimental model for studying the evolution of Apicomplexa [3], particularly the transition from phototrophic algae to obligate parasites of animals.

Temperature is one of the most obvious axis along which such divergence might arise (most easy to reproduce), and it is also the most pressing, given the accelerating warming of the oceans in which these algae live. To address this, we compared two isolates separated by roughly 10° of latitude (∼1100 km), obtained from Sydney Harbour and the Southern Great Barrier Reef (Fig 1A). These sites differ markedly in their thermal regimes (Fig 1B and 1C): the southern GBR is substantially warmer, whereas Sydney Harbour is both cooler and more variable in temperature. Both sites have experienced increasingly frequent and intense warm anomalies since 2015 (Fig 1B), making the question of how each population responds to elevated temperature directly relevant to their persistence under ongoing climate change.

### Strain divergence dominates transcriptomic variation

Despite the morphological and ultrastructural similarity of the two isolates, the isolates differ in their transcriptomic and lipidomic responses to tropical temperatures. Principal component analysis identified strain identity as the primary axis of transcriptomic variation, exceeding the temperature effect by more than an order of magnitude (Fig 3C). This indicates that the two isolates have diverged substantially in baseline gene regulation, independent of the immediate thermal condition. Beyond a stable core transcriptome expressed across all four conditions, OTI expressed more genes than SH at both temperatures, including more than three thousand genes never detected in SH under either condition (Fig 3A and 3B). A further set of genes expressed by both strains at 20°C was specifically lost from the SH profile at 26°C while remaining expressed in OTI. Broader expressed-gene diversity in OTI was therefore not an artifact of a single condition but a consistent feature of the strain, and the gap between the two widened specifically under heat (Fig 3B). Increased temperature added a smaller set of differences on top of this pattern: about a thousand genes differed between the strains only at 26°C, where they were upregulated in OTI, and a smaller group differed only at 20°C, where they were upregulated in SH (Fig 4E). Genes that switched direction, upregulated in one strain at 20°C and in the other at 26°C, were rare (Fig 4E). Temperature therefore determined a magnitude of differences between the strains, not which strain shows gene upregulation. Similar difference between conspecific populations originating from different thermal conditions have been described in reef corals, where the transcriptional profile of a heat-tolerant population differed from that of more sensitive neighbor in its response to heat [40].

### Strains display different transcriptional responses to increased temperature

Increasing temperature from 20°C to 26°C produced markedly different responses in the two strains. In SH, far more genes were suppressed than activated, whereas in OTI up- and downregulation were more evenly balanced (Fig 5A and 5B). Downregulation-dominated responses to heat have been reported elsewhere, although the proportion varies: about 62% of differentially expressed genes were downregulated in the green alga *Scenedesmus* [41] and about 72% in the bacterium *Mesorhizobium loti* under heat shock [42]. The proportion of downregulated genes in SH is higher than in either case (Fig 5A). In other lineages the same cue produces a largely upregulated transcriptome in the rhodophyte *Bangia fuscopurpurea* [43], or an up-skewed one in which only specific functions, such as translation and photosynthesis, are suppressed [44]. Broad suppression has been interpreted both as a marker of damage and as an energy-saving strategy [42,45], so on its own it does not distinguish stress from tolerance. The same membrane-to-storage lipid conversion we documented in *C. velia* under heat (Fig 8) has previously been elicited in this strain by a completely different stressor, nitrogen depletion, suggesting a general SH stress program rather than a temperature-specific one [46].

However, we observed that the growth rate and final OD600 were higher at 26°C than at 20°C in both strains (Fig 2A and 2C). Moreover, zoospores, a motile stage of the alga, appeared only at the higher temperature (Fig 2B). This argues against 26°C being heat stress condition for SH. Heat-induced zoosporogenesis (gametogenesis) is a strategy documented for other algae, for example the green alga *Volvox carteri*, where heat shock specifically, and no other stressors tested, triggers a shift toward sexual reproduction, interpreted as an anticipatory response to deteriorating habitat conditions [47]. Even more similarly, because *C. velia* is asexual, a strain-specific pattern has been reported for the rhodophyte genus *Bangia*, in which heat-triggered asexual spore release coincides with acquired heat tolerance in one strain, whereas closely related strains neither release spores under heat nor acquire tolerance from it [48]. The reverse effect is also documented in cereals, where heat and drought stress damage, rather than promote reproductive output, reducing pollen viability and fertilization success [49].

Some metabolic pathways can be selectively activated. For example, SH’s photosynthesis-associated transcripts were upregulated under heat rather than downregulated[43,50], and a similarly selective, non-uniform response, with some components suppressed, others maintained or induced, has been described in the dinoflagellate *Prorocentrum cordatum* under heat [44]. What is evident, however, is that SH’s response to heat differs fundamentally from OTI’s at every level we measured, transcriptome (Fig 3C), lipidome (Fig 8), and functional gene-set direction (Fig 6, Fig 7; discussed further below), and it is this divergence between strains in particular, rather than the interpretation attached to either one, that is the central finding of this study.

### The strains activate opposite transcriptional programs in response to the same thermal cue

Gene ontology (GO) enrichment (Fig 6) and gene-level analyses (Fig 7) show that the two strains differ not only in the intensity of their transcriptional response, but also in the selective activation of genes involved in functions spanning core cellular processes. Some of these functions are associated with thermal tolerance in other algae. In Symbiodiniaceae, the more heat-tolerant species maintained synthesis and repair of biomolecules under heat stress, whereas the more sensitive species showed a decline in anabolic processes [45]. The cilium/microtubule-motility system is the clearest example here (Fig 7D): motility gene expression was comparable between *C. velia* strains at 20°C, but markedly higher in OTI at 26°C (Fig 7D, rings 1–2), with cilium- and microtubule-related functions among the most strongly elevated OTI categories in the 26°C strain comparison (Fig 6B).

The strain difference become particularly pronounced at the higher temperature because SH shows inhibited expression, not because OTI gains expression. The motility genes involved in microtubule- and cilium-related functions were the most strongly downregulated (Fig 6D) in SH at the higher temperature (Fig 7D, ring 3), while these expression of these genes remained unchanged in OTI under the same conditions (Fig 7D, ring 4). Because the comparison of strains at 26°C and comparison of SH at both temperatures share the same SH 26°C measurement, these are not independently obtained data. However, they demonstrate the divergence in motility gene downregulation in SH at different temperatures, rather than activation of gene expression in OTI (Fig 6B, 6D and 7D). A comparable downregulation of cytoskeletal and microtubule-associated transcripts under heat has been reported in the dinoflagellate *Prorocentrum cordatum* [44], so this phenomenon is very likely general, at least among alveolates.

An even greater divergence appeared in the lipid-related gene set (Fig 7C): most lipid-related genes were upregulated in OTI and downregulated in SH at both temperatures, however, the change seemed coordinated in OTI. A probable S-acyltransferase (Cvel_8830) and a lysophospholipid acyltransferase LPEAT1 (Cvel_12962) were upregulated in OTI compared with SH at 26°C.A phospholipid-transporting ATPase (Cvel_8341) showed the same pattern. This coordinated suppression of lipid-remodeling machinery in SH is the transcriptional counterpart of the membrane-to-storage lipid shift documented for this strain at the lipidome level (Fig 8), and is consistent with heat-induced membrane-lipid remodeling described in other microalgae [51,52]. In Symbiodiniaceae, heat-sensitive taxa extensively remodel their membranes, whereas thermotolerant taxa preserve membrane lipid composition [53], and shifts in betaine lipid and galactolipid ratios track heat tolerance across species [54]. SH’s transricptomic pattern of lipid-related genes is similar to the former, and OTI’s to the latter.

The ribosome/translation gene set (Fig 7B) showed the opposite transcription pattern, with most genes downregulated in OTI compared with SH at both temperatures, and translation and biosynthesis functions dominating the SH-elevated categories in the 26°C strain comparison (Fig 6B). This should not be interpreted as active induction of gene expression upon heating, as it reflects a between-strain difference at a single temperature, along with differential transcription, mostly downregulation, in SH at different temperatures (Fig 7B). This suggests that the two isolates of *C. velia* respond to different temperatures using distinct metabolic programs. It should be noted that SH is the only *C. velia* isolate available in public culture collections, and this isolate has therefore been used in most experiments [10]. At the same time, the cultivation temperatures recommended by the NCMA culture collection are 24-26°C. Across marine phytoplankton, the thermal optimum of a strain tracks the temperature at its site of isolation [55], and in the coccolithophore *Gephyrocapsa huxleyi*, strains from different thermal regimes use distinct thermal response mechanisms, so no single isolate captures the species’ response [56]. Our data suggest the same holds for *C. velia*.

### Lipidomic shows a conserved thermal response

Much of what the two isolates did at 26 °C, they did similarly. Both reduced their MGDG:DGDG ratio by more than 80 %, and both depleted MGDG and the anionic photosystem lipids SQDG and PG, with no strain × temperature interaction in any of these comparisons. The direction of this change is the one predicted from membrane biophysics. MGDG is conical and non-bilayer-forming, whereas DGDG is cylindrical and bilayer-stabilizing, so a fall in their ratio shifts the membrane toward a lamellar conformation [57]. In Arabidopsis, DGDG-deficient thylakoids are specifically compromised at elevated temperature, and the capacity of plants to acquire thermal tolerance correlates with an increase in DGDG and in the DGDG:MGDG ratio rather than with heat-shock protein accumulation [58]. Comparable galactolipid remodeling, with loss of the most unsaturated MGDG species and retention of more saturated DGDG, has been documented in heat-treated tomato [59].

Both isolates also withdrew eicosapentaenoic acid from membrane lipids and recovered it in the storage pool, and the interaction was significant only on the membrane side — that is, the two isolates removed different amounts, but what they removed ended up in the same destination. This route has been established in other algae. In Chlamydomonas reinhardtii, heat-induced triacylglycerol carries the acyl signature of the membrane lipids that disappear, which distinguishes direct conversion from de novo synthesis, and candidate lipases and acyltransferases have been identified [52]. Betaine lipids serve as acyl donors for this transfer in *Phaeodactylum tricornutum* [60], and the same coupling between plastid membrane recycling and neutral lipid formation operates across diatoms [61]. In Coccomyxa, isotope labeling showed that heat-induced oil accumulation arises from redistribution of carbon flux rather than from increased fatty acid synthesis [62].

This conserved component is worth stating plainly, because it is not what the lipid-class data alone suggested. OTI does remodel its membrane at 26 °C; it is not thermally inert.

### OTI membranes are constitutively less unsaturated

The temperature-independent difference is the more interesting one. At 20 °C, OTI membrane lipids carried 1.92 ± 0.07 double bonds per acyl chain compared with 2.37 ± 0.04 in SH, with polyunsaturated chains at 38.9 % versus 49.5 %. The separation persisted at 26 °C with no interaction term and was absent from the storage pool at 20 °C. OTI therefore maintains a membrane that is already closer to a heat-tolerant configuration, regardless of the temperature at which it is grown.

This is precisely the relationship proposed for coral symbionts. Tchernov and colleagues showed that the critical threshold temperature separating thermally tolerant from thermally sensitive *Symbiodinium* is set by the degree of saturation of their thylakoid lipids, and argued that membrane lipid composition is diagnostic of bleaching susceptibility [63]. The hypothesis has not gone unchallenged: a later comparison of Cladocopium C1 and Symbiodinium A1 found no straightforward relationship between fatty acid saturation, membrane stability, and thermal tolerance [64], and subsequent lipidomic work has implicated betaine and galactolipid class ratios alongside, or instead of, bulk unsaturation [53,54,65]. Our data cannot settle that debate, but they add an unusual case to it. The comparison here is not between distantly related symbiont lineages with different evolutionary histories, but between two isolates of a single described species with closely similar genomic backgrounds, which removes much of the phylogenetic confounding that complicates the Symbiodiniaceae comparisons.

The underlying logic is older than the coral literature. Homeoviscous adaptation predicts that membrane acyl unsaturation is tuned to the prevailing temperature so as to keep membrane physical properties approximately constant [66,67]. A lineage from a habitat that is on average 3.4 °C warmer would be expected to sit at a lower unsaturation set point, which is what we observe.

### The isolates differ in where new monounsaturated chains are placed

The clearest strain-specific signal in the dataset is the fate of oleic acid. Both isolates contained more 18:1n-9 at 26 °C than at 20 °C, so both were synthesizing it. However, OTI incorporated it into membrane lipids, whereas SH routed it into storage, with a significant strain × temperature interaction in both pools. This pattern matches a mechanism described in fission yeast. Péter and colleagues proposed that heat-induced triacylglycerol synthesis contributes to membrane rigidization by accommodating unsaturated acyl chains withdrawn from structural lipids, allowing them to be replaced by newly synthesized saturated ones [68]; cells that cannot synthesize triacylglycerol are correspondingly less able to adjust their membrane physical properties under heat [69]. The mechanism has two parts: the removal of unsaturated chains and their replacement. Our data suggest that the two isolates differ in whether the second part occurs. Both withdrew polyunsaturated chains from the membrane. Only OTI replaced them, and its membrane pool expanded. In SH, the withdrawn chains were not replaced, the newly synthesized monounsaturated chains went to storage instead, and the membrane pool contracted by roughly half. Framed this way, SH is not failing to mount a response; It is executing only half of one. That interpretation is consistent with the transcriptomic layer, in which fifteen of eighteen lipid-related genes examined were downregulated in SH at 26 °C, whereas OTI showed no coordinated change, although we emphasize that the correspondence between the two layers is a correlation across two clonal lineages and does not establish that the transcriptional change causes the lipid phenotype.

## Conclusions

A previous comparison of two *C. velia* isolates, the Sydney strain and the Mdig3 strain isolated from *Montipora digitata* at Magnetic Island in the central Great Barrier Reef [14], found them highly similar in sequence and concluded that results obtained with one could be generalized to the other [70]. The same study identified two HSP20 isoforms in both *C. velia* strains, absent from the Symbiodiniaceae examined, and proposed that these small heat shock proteins may underlie the relatively high thermal tolerance of *C. velia* [70]. Our two isolates, however, respond to the same increase in temperature in markedly different ways. In SH, an increase from 20°C to 26°C caused a dominant suppression of motility, cytoskeletal and lipid-remodeling functions, together with a contraction of the membrane lipid pool relative to storage lipids. Its lipidome also changed, but newly synthesized oleic acid was directed into membrane lipids, and its membranes were less unsaturated (requiring less remodeling) than those of SH at both temperatures. Because unsaturated acyl chains keep membranes fluid, and warming makes membranes more fluid still, a membrane that is already less unsaturated is consistent with a lipid composition pre-adjusted to warmer water and. The HSP20 genes proposed as a basis for *C. velia* thermal tolerance were induced at 26°C in both isolates, so both registered the change in temperature, yet their overall responses still diverged. Isolates of the same species can therefore differ substantially in how they respond to warming, and results obtained with *C. velia* isolates should be carefully interpreted taking their different thermotolerance into consideration (at least for OTI strain of *Chromera velia*). As marine heatwaves intensify and warm anomalies become more frequent at both isolation sites, the higher thermal stability documented in OTI might play an important role in stabilizing the coral environment.

## Supporting information

https://zenodo.org/uploads/23415855?token=eyJhbGciOiJIUzUxMiJ9.eyJpZCI6IjA2MTkxY2UzLTE0NzEtNGIyYi1hYTk5LWU3OTQ1M2ZkYjgxOCIsImRhdGEiOnt9LCJyYW5kb20iOiI

## Acknowledgments

We thank Dr. Ansgar Gruber for insightful discussions regarding this project. We also thank Antonín Střížek and Pavel Hrouzek for cooperation and for using equipment at Algatech during pilot experiments. We also thank the Electron Microscopy Facility at the Institute of Parasitology, Biology Centre CAS, for imaging our samples.

## Funding

This work is supported by Czech Science Foundation grant no 25-15611S. The funders had no role in study design, data collection and analysis, decision to publish, or preparation of the manuscript.

## Competing interests

The authors declare that there was no potential conflict of interest.

## Author’s Contribution

**Conceptualization:** Ayush Sharma, Aleš Horák, Miroslav Oborník.

**Formal analysis:** Ayush Sharma, Aleš Tomčala, Tomáš Skalický, Aleš Horák.

**Funding acquisition:** Miroslav Oborník.

**Investigation:** Ayush Sharma, Aleš Tomčala, Tomáš Skalický, Dorsaf Ennaceur, Jitka Richtová, Vijaya Geetha Gonepogu.

**Methodology:** Ayush Sharma, Aleš Tomčala, Tomáš Skalický, Aleš Horák, Miroslav Oborník.

**Project administration:** Miroslav Oborník.

**Resources:** Jan Šlapeta, Miroslav Oborník.

**Supervision:** Miroslav Oborník.

**Writing-original draft:** Ayush Sharma, Aleš Tomčala, Miroslav Oborník.

**Writing-review & editing:** Ayush Sharma, Aleš Tomčala, Tomáš Skalický, Aleš Horák, Jan Šlapeta, Miroslav Oborník.

## Data availability statement

Raw RNA-seq reads have been deposited in the NCBI Sequence Read Archive under BioProject PRJNA1519432. The analysis code, processed data, the gene count matrix, all numerical values underlying the figures, and annotated Excel workbooks listing the differentially expressed genes for each comparison and direction (including GO enrichment results) are archived at Zenodo and can be accessed with link <u>Supplementary data</u>.

## Supplementary legends

**S1 Fig. Cell morphology of the OTI and SH strains of *Chromera velia* visualized by scanning and transmission electron microscopy**

(**A**, **B**) Scanning electron micrographs (SEM) of the tropical OTI strain, isolated from One Tree Island (southern Great Barrier Reef), showing (A) an individual coccoid vegetative cell and (B) a chain of adjacent vegetative cells at lower magnification. (**C**, **D**) Scanning electron micrographs of the SH strain, isolated from Sydney Harbour, showing (C) an individual vegetative cell and (D) a cluster of multiple coccoid cells. (E, F) Transmission electron micrographs (TEM) of representative cross-sections of (E) the OTI strain and (F) the SH strain, showing the major intracellular compartments including the nucleus, plastid with stacked thylakoid membranes, and storage lipid bodies.

**S2 Fig. Comparison of cell size between the SH and OTI strains of *Chromera velia*.**

Bar plot showing the mean maximum cell diameter (Dmax, μm) of the SH (yellow) and OTI (green) strains, quantified from scanning electron micrographs. Individual cell measurements are provided in S1 Table (SH: n = 23 cells, mean = 5.98 ± 0.64 μm; OTI: n = 21 cells, mean = 5.91 ± 0.72 μm); error bars represent standard deviation across cells. Cell size did not differ meaningfully between the two strains. Representative micrographs (right) illustrate the coccoid morphology and cells scored for Dmax; white lines indicate the axis along which the maximum diameter was measured, and numbers correspond to cell IDs in Supplementary Table 1.

**S3 Fig. Phylogenetic tree.**

18S rDNA phylogenetic tree placing the OTI isolate among Apicomplexa and related alveolates.

**S1 Table. Isolates of *Chromera velia*.**

Table showing *Chromera velia* isolates, isolation location, associated coral, isolation temperature recorded and reference articles

**S2 Table.** Individual cell measurements used to build **S2 Fig.** (SH: n = 23 cells, mean = 5.98 ± 0.64 μm; OTI: n = 21 cells, mean = 5.91 ± 0.72 μm);

## Notes

https://zenodo.org/uploads/23415855?token=eyJhbGciOiJIUzUxMiJ9.eyJpZCI6IjA2MTkxY2UzLTE0NzEtNGIyYi1hYTk5LWU3OTQ1M2ZkYjgxOCIsImRhdGEiOnt9LCJyYW5kb20iOiI4ZGJhYTdiNjk4YzI5NmUzMGY0OThkMDhhNGQ0MDBmZSJ9.dM52bwZ9P8WTc3E5N2ZfCqLaNNvoCs13k_vHgKvZBNnjSRUkB2y_F6Wvwh5SXl4uMiXBX7__F5Eu2v6mYlh5RA

### Competing Interest Statement

The authors have declared no competing interest.

## Reference

1. Votýpka J, Modrý D, Oborník M, Šlapeta J, Lukeš J. Apicomplexa. Handbook of the Protists. Cham: Springer International Publishing; 2016. pp. 1–58. doi:10.1007/978-3-319-32669-6_20-1

2. McFadden GI, Reith ME, Munholland J, Lang-Unnasch N. Plastid in human parasites. Nature. 1996;381: 482. doi:10.1002/bies.950191114

3. Cavalier-Smith T. Kingdom Chromista and its eight phyla: a new synthesis emphasising periplastid protein targeting, cytoskeletal and periplastid evolution, and ancient divergences. Protoplasma. 2018;255: 297–357. doi:10.1007/s00709-017-1147-3

4. Oborník M. Endosymbiotic evolution of algae, secondary heterotrophy and parasitism. Biomolecules. 2019;9: 266. doi:10.3390/biom9070266

5. Oborník M. Photoparasitism as an Intermediate State in the Evolution of Apicomplexan Parasites. Trends Parasitol. 2020;36: 727–734. doi:10.1016/j.pt.2020.06.002

6. Zhu G, Marchewka MJ, Keithly JS. Cryptosporidium parvum appears to lack a plastid genome. Microbiology (N Y). 2000;146: 315–321. doi:10.1099/00221287-146-2-315

7. Toso MA, Omoto CK. Gregarina niphandrodes may lack both a plastid genome and organelle. J Eukaryot Microbiol. 2007;54: 66–72. doi:10.1111/j.1550-7408.2006.00229.x

8. Jacko-Reynolds VKL, Kwong WK, Livingston SJ, Trznadel M, Bonacolta AM, Lax G, et al. Phylogenomics of coral-infecting corallicolids reveal multiple independent losses of chlorophyll biosynthesis in apicomplexan parasites. Current Biology. 2025;35: 1156–1163.e4. doi:10.1016/j.cub.2025.01.028

9. Janouškovec J, Paskerova GG, Miroliubova TS, Mikhailov K V, Birley T, Aleoshin V V, et al. Apicomplexan-like parasites are polyphyletic and widely but selectively dependent on cryptic plastid organelles. Elife. 2019;8. doi:10.7554/eLife.49662

10. Moore RB, Oborník M, Janouškovec J, Chrudimský T, Vancová M, Green DH, et al. A photosynthetic alveolate closely related to apicomplexan parasites. Nature. 2008;451: 959–963. doi:10.1038/nature06635

11. Oborník M, Vancová M, Lai D-H, Janouškovec J, Keeling PJ, Lukeš J. Morphology and Ultrastructure of Multiple Life Cycle Stages of the Photosynthetic Relative of Apicomplexa, Chromera velia. Protist. 2011;162: 115–130. doi:10.1016/j.protis.2010.02.004

12. Oborník M, Modrý D, Lukeš M, Černotíková-Stříbrná E, Cihlář J, Tesařová M, et al. Morphology, Ultrastructure and Life Cycle of Vitrella brassicaformis n. sp., n. gen., a Novel Chromerid from the Great Barrier Reef. Protist. 2012;163: 306–323. doi:10.1016/j.protis.2011.09.001

13. Füssy Z, Oborník M. Chapter Six - Chromerids and Their Plastids. In: Hirakawa Y, editor. Advances in Botanical Research. Academic Press; 2017. pp. 187–218. 10.1016/bs.abr.2017.07.001

14. Cumbo VR, Baird AH, Moore RB, Negri AP, Neilan BA, Salih A, et al. Chromera velia is Endosymbiotic in Larvae of the Reef Corals Acropora digitifera and A. tenuis. Protist. 2013;164: 237–244. doi:10.1016/j.protis.2012.08.003

15. Mohamed AR, Cumbo VR, Harii S, Shinzato C, Chan CX, Ragan MA, et al. Deciphering the nature of the coral– *Chromera* association. ISME J. 2018;12: 776–790. doi:10.1038/s41396-017-0005-9

16. Füssy Z, Masařová P, Kručinská J, Esson HJ, Oborník M. Budding of the alveolate alga Vitrella brassicaformis resembles sexual and asexual processes in apicomplexan parasites. Protist. 2017;168: 80–91.

17. Janouškovec J, Horák A, Barott KL, Rohwer FL, Keeling PJ. Environmental distribution of coral-associated relatives of apicomplexan parasites. ISME J. 2013;7: 444.

18. Bonacolta AM, Keeling PJ. Modern microbialites harbor an undescribed diversity of chromerid algae. Environ Microbiome. 2026;21: 25. doi:10.1186/s40793-026-00852-4

19. Weatherby K, Murray S, Carter D, Šlapeta J. Surface and Flagella Morphology of the Motile Form of Chromera velia Revealed by Field-Emission Scanning Electron Microscopy. Protist. 2011;162: 142–153. doi:10.1016/j.protis.2010.02.003

20. Morin-Adeline V, Foster C, Šlapeta J. Identification of Chromera velia by fluorescence in situ hybridization. FEMS Microbiol Lett. 2012;328: 144–149. doi:10.1111/j.1574-6968.2011.02489.x

21. Cumbo VR. Antimicrobial compounds in the scleractinian corals Montipora digitata and Montipora tortuosa. 2005.

22. Visser P, Bintoudi E, Boschker E, Frade P, van Bleijswijk J, Matthijs H. Newly discovered coral endosymbiont Chromera is more thermotolerant than Symbiodinium. 12th International Coral Reef Symposium Abstracts (Cairns). 2012.

23. Richtová J, Ennaceur D, Oborník M. Separating Chromera velia Zoospores From Culture and Estimating Their Average Motility Speed and Lifespan. Bio Protoc. 2026;16. doi:10.21769/BioProtoc.5756

24. Bolger AM, Lohse M, Usadel B. Trimmomatic: a flexible trimmer for Illumina sequence data. Bioinformatics. 2014;30: 2114–2120. doi:10.1093/bioinformatics/btu170

25. Dobin A, Davis CA, Schlesinger F, Drenkow J, Zaleski C, Jha S, et al. STAR: ultrafast universal RNA-seq aligner. Bioinformatics. 2013;29: 15–21.

26. Robinson MD, McCarthy DJ, Smyth GK. edgeR: a Bioconductor package for differential expression analysis of digital gene expression data. bioinformatics. 2010;26: 139–140.

27. Love MI, Huber W, Anders S. Moderated estimation of fold change and dispersion for RNA-seq data with DESeq2. Genome Biol. 2014;15: 550. doi:10.1186/s13059-014-0550-8

28. Alvarez-Jarreta J, Amos B, Aurrecoechea C, Bah S, Barba M, Barreto A, et al. VEuPathDB: the eukaryotic pathogen, vector and host bioinformatics resource center in 2023. Nucleic Acids Res. 2024;52: D808–D816. doi:10.1093/nar/gkad1003

29. Folch J, Lees M, Stanley GHS. A SIMPLE METHOD FOR THE ISOLATION AND PURIFICATION OF TOTAL LIPIDES FROM ANIMAL TISSUES. Journal of Biological Chemistry. 1957;226: 497–509. doi:10.1016/S0021-9258(18)64849-5

30. Tomčala A, Kyselová V, Schneedorferová I, Opekarová I, Moos M, Urajová P, et al. Separation and identification of lipids in the photosynthetic cousins of Apicomplexa *Chromera velia* and *Vitrella brassicaformis*. J Sep Sci. 2017;40: 3402–3413. doi:10.1002/jssc.201700171

31. Boyer TP, Antonov JI, Baranova OK, Garcia HE, Johnson DR, Mishonov A V, et al. World ocean database 2013. 2013.

32. Hobday AJ, Pecl GT. Identification of global marine hotspots: sentinels for change and vanguards for adaptation action. Rev Fish Biol Fish. 2014;24: 415–425. doi:10.1007/s11160-013-9326-6

33. Lough JM, Anderson KD, Hughes TP. Increasing thermal stress for tropical coral reefs: 1871–2017. Sci Rep. 2018;8: 6079. doi:10.1038/s41598-018-24530-9

34. Howells EJ, Beltran VH, Larsen NW, Bay LK, Willis BL, van Oppen MJH. Coral thermal tolerance shaped by local adaptation of photosymbionts. Nat Clim Chang. 2012;2: 116–120. doi:10.1038/nclimate1330

35. Yuan M-H, Lin K-T, Pan S-Y, Yang C-K. Exploring coral reef benefits: A systematic SEEA-driven review. Science of The Total Environment. 2024;950: 175237. doi:10.1016/j.scitotenv.2024.175237

36. Wiedenmann J, D’Angelo C, Mardones ML, Moore S, Benkwitt CE, Graham NAJ, et al. Reef-building corals farm and feed on their photosynthetic symbionts. Nature. 2023;620: 1018–1024. doi:10.1038/s41586-023-06442-5

37. Janouškovec J, Horák A, Oborník M, Lukeš J, Keeling PJ. A common red algal origin of the apicomplexan, dinoflagellate, and heterokont plastids. Proceedings of the National Academy of Sciences. 2010;107: 10949–10954. doi:10.1073/pnas.1003335107

38. Flegontov P, Michálek J, Janouškovec J, Lai D-H, Jirků M, Hajdušková E, et al. Divergent mitochondrial respiratory chains in phototrophic relatives of apicomplexan parasites. Mol Biol Evol. 2015;32: 1115–1131. doi:10.1093/molbev/msv021

39. Woo YH, Ansari H, Otto TD, Klinger CM, Kolisko M, Michálek J, et al. Chromerid genomes reveal the evolutionary path from photosynthetic algae to obligate intracellular parasites. Elife. 2015;4. doi:10.7554/eLife.06974

40. Barshis DJ, Ladner JT, Oliver TA, Seneca FO, Traylor-Knowles N, Palumbi SR. Genomic basis for coral resilience to climate change. Proceedings of the National Academy of Sciences. 2013;110: 1387–1392.

41. Calhoun S, Bell TAS, Dahlin LR, Kunde Y, LaButti K, Louie KB, et al. A multi-omic characterization of temperature stress in a halotolerant Scenedesmus strain for algal biotechnology. Commun Biol. 2021;4: 333. doi:10.1038/s42003-021-01859-y

42. Alexandre A, Laranjo M, Oliveira S. Global Transcriptional Response to Heat Shock of the Legume Symbiont Mesorhizobium loti MAFF303099 Comprises Extensive Gene Downregulation. DNA Research. 2014;21: 195–206. doi:10.1093/dnares/dst050

43. Zhao M, Zheng H, Chen Z, Chen W. Physiological and Biochemical Responses and Transcriptome Analysis of Bangia fuscopurpurea (Rhodophyta) Under High-Temperature Stress. Curr Issues Mol Biol. 2025;47. doi:10.3390/cimb47070484

44. Dougan KE, Deng Z-L, Wöhlbrand L, Reuse C, Bunk B, Chen Y, et al. Multi-omics analysis reveals the molecular response to heat stress in a “red tide” dinoflagellate. Genome Biol. 2023;24: 265. doi:10.1186/s13059-023-03107-4

45. Xiang T, Peak SL, Huitt EC, Grossman AR. Distinct transcriptomic strategies underlie differential heat tolerance in Symbiodiniaceae symbionts. ISME J. 2025;19. doi:10.1093/ismejo/wraf268

46. Tomčala A, Michálek J, Schneedorferová I, Füssy Z, Gruber A, Vancová M, et al. Fatty Acid Biosynthesis in Chromerids. Biomolecules. 2020;10: 1102. doi:10.3390/biom10081102

47. Kirk DL, Kirk MM. Heat shock elicits production of sexual inducer in Volvox. Science (1979). 1986;231: 51–54.

48. Mikami K. Research on the Regulatory Mechanism of Algae Reproduction under Abiotic Stress Conditions. Plants. 2022;11: 525. doi:10.3390/plants11040525

49. Barnabás B, Jäger K, Fehér A. The effect of drought and heat stress on reproductive processes in cereals. Plant Cell Environ. 2008;31: 11–38.

50. Gierz SL, Forêt S, Leggat W. Transcriptomic analysis of thermally stressed Symbiodinium reveals differential expression of stress and metabolism genes. Front Plant Sci. 2017;8. doi:10.3389/fpls.2017.00271

51. Hemme D, Veyel D, Mühlhaus T, Sommer F, Jüppner J, Unger A-K, et al. Systems-Wide Analysis of Acclimation Responses to Long-Term Heat Stress and Recovery in the Photosynthetic Model Organism *Chlamydomonas reinhardtii*. Plant Cell. 2014;26: 4270–4297. doi:10.1105/tpc.114.130997

52. Légeret B, Schulz Raffelt M, Nguyen HM, Auroy P, Beisson F, Peltier G, et al. Lipidomic and transcriptomic analyses of *Chlamydomonas reinhardtii* under heat stress unveil a direct route for the conversion of membrane lipids into storage lipids. Plant Cell Environ. 2016;39: 834–847. doi:10.1111/pce.12656

53. T. Botana M, Chaves Filho AB, Inague A, Z. Güth A, Saldanha Corrêa F, Müller MN, et al. Thermal plasticity of coral reef symbionts is linked to major alterations in their lipidome composition. Limnol Oceanogr. 2022;67: 1456–1469. doi:10.1002/lno.12094

54. Rosset S, Koster G, Brandsma J, Hunt AN, Postle AD, D’Angelo C. Lipidome analysis of Symbiodiniaceae reveals possible mechanisms of heat stress tolerance in reef coral symbionts. Coral Reefs. 2019;38: 1241–1253. doi:10.1007/s00338-019-01865-x

55. Thomas MK, Kremer CT, Klausmeier CA, Litchman E. A Global Pattern of Thermal Adaptation in Marine Phytoplankton. Science (1979). 2012;338: 1085–1088. doi:10.1126/science.1224836

56. Krinos AI, Shapiro SK, Li W, Haley ST, Dyhrman ST, Dutkiewicz S, et al. Intraspecific Diversity in Thermal Performance Determines Phytoplankton Ecological Niche. Ecol Lett. 2025;28. doi:10.1111/ele.70055

57. Yu C, Lin Y, Li H. Increased ratio of galactolipid MGDG : DGDG induces jasmonic acid overproduction and changes chloroplast shape. New Phytologist. 2020;228: 1327–1335. doi:10.1111/nph.16766

58. Krumova SB, Laptenok SP, Kovács L, Tóth T, van Hoek A, Garab G, et al. Digalactosyl-diacylglycerol-deficiency lowers the thermal stability of thylakoid membranes. Photosynth Res. 2010;105: 229–242. doi:10.1007/s11120-010-9581-5

59. Spicher L, Glauser G, Kessler F. Lipid Antioxidant and Galactolipid Remodeling under Temperature Stress in Tomato Plants. Front Plant Sci. 2016;7. doi:10.3389/fpls.2016.00167

60. Popko J, Herrfurth C, Feussner K, Ischebeck T, Iven T, Haslam R, et al. Metabolome Analysis Reveals Betaine Lipids as Major Source for Triglyceride Formation, and the Accumulation of Sedoheptulose during Nitrogen-Starvation of Phaeodactylum tricornutum. PLoS One. 2016;11: e0164673. doi:10.1371/journal.pone.0164673

61. Murison V, Hérault J, Côme M, Guinio S, Lebon A, Chamot C, et al. Comparison of two Phaeodactylum tricornutum ecotypes under nitrogen starvation and resupply reveals distinct lipid accumulation strategies but a common degradation process. Front Plant Sci. 2023;14. doi:10.3389/fpls.2023.1257500

62. Allen JW, Tevatia R, Demirel Y, DiRusso CC, Black PN. Induction of oil accumulation by heat stress is metabolically distinct from N stress in the green microalgae Coccomyxa subellipsoidea C169. PLoS One. 2018;13: e0204505.

63. Tchernov D, Gorbunov MY, de Vargas C, Narayan Yadav S, Milligan AJ, Häggblom M, et al. Membrane lipids of symbiotic algae are diagnostic of sensitivity to thermal bleaching in corals. Proceedings of the National Academy of Sciences. 2004;101: 13531– 13535. doi:10.1073/pnas.0402907101

64. Díaz-Almeyda E, Thomé PE, El Hafidi M, Iglesias-Prieto R. Differential stability of photosynthetic membranes and fatty acid composition at elevated temperature in Symbiodinium. Coral Reefs. 2011;30: 217–225.

65. Tortorelli G, Rosset SL, Sullivan CES, Woo S, Johnston EC, Walker NS, et al. Heat-induced stress modulates cell surface glycans and membrane lipids of coral symbionts. ISME J. 2025;19. doi:10.1093/ismejo/wraf073

66. Sinensky M. Homeoviscous Adaptation—A Homeostatic Process that Regulates the Viscosity of Membrane Lipids in *Escherichia coli*. Proceedings of the National Academy of Sciences. 1974;71: 522–525. doi:10.1073/pnas.71.2.522

67. Hazel JR. Thermal Adaptation in Biological Membranes: Is Homeoviscous Adaptation the Explanation? Annu Rev Physiol. 1995;57: 19–42. doi:10.1146/annurev.ph.57.030195.000315

68. Péter M, Glatz A, Gudmann P, Gombos I, Török Z, Horváth I, et al. Metabolic crosstalk between membrane and storage lipids facilitates heat stress management in Schizosaccharomyces pombe. PLoS One. 2017;12: e0173739. doi:10.1371/journal.pone.0173739

69. Gudmann P, Gombos I, Péter M, Balogh G, Török Z, Vígh L, et al. Mild Heat Stress Alters the Physical State and Structure of Membranes in Triacylglycerol-Deficient Fission Yeast, Schizosaccharomyces pombe. Cells. 2024;13: 1543. doi:10.3390/cells13181543

70. Mohamed AR, Chan CX, Ragan MA, Zhang J, Cooke I, Ball EE, et al. Comparative transcriptomic analyses of *Chromera* and Symbiodiniaceae. Environ Microbiol Rep. 2020;12: 435–443. doi:10.1111/1758-2229.12859

