## Supplementary material for "Transcriptomic and lipidomic profiles reveal higher thermal adaptability in a tropical versus a temperate isolate of *Chromera velia*": https://zenodo.org/uploads/23415855?token=eyJhbGciOiJIUzUxMiJ9.eyJpZCI6IjA2MTkxY2UzLTE0NzEtNGIyYi1hYTk5LWU3OTQ1M2ZkYjgxOCIsImRhdGEiOnt9LCJyYW5kb20iOiI

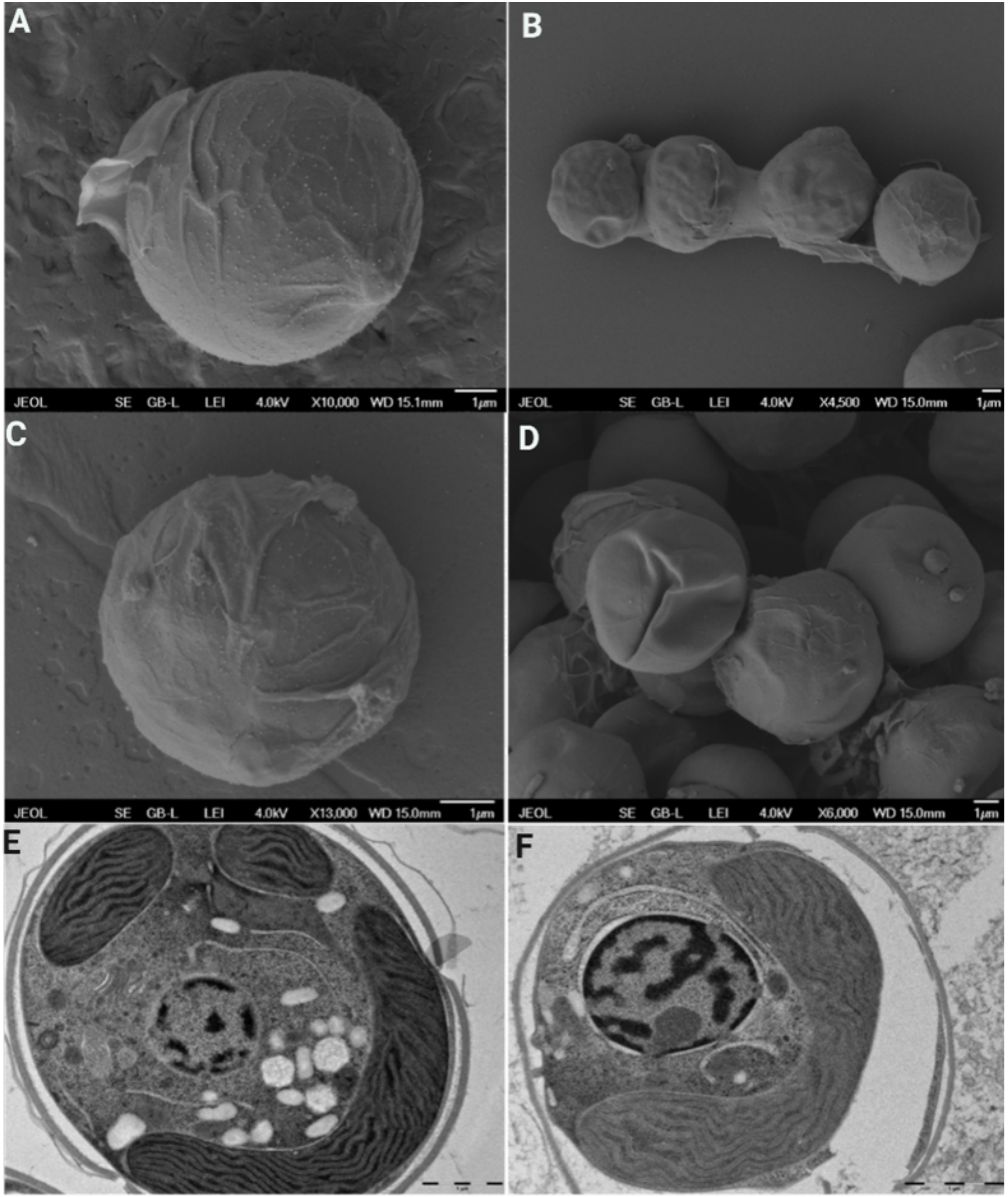

**S1 Fig. Cell morphology of the OTI and SH strains of *Chromera velia* visualized by scanning and transmission electron microscopy.** (**A**, **B**) Scanning electron micrographs (SEM) of the tropical OTI strain, isolated from One Tree Island (southern Great Barrier Reef), showing (A) an individual coccoid vegetative cell and (B) a chain of adjacent vegetative cells at lower magnification. (**C**, **D**) Scanning electron micrographs of the SH strain, isolated from Sydney Harbour, showing (C) an individual vegetative cell and (D) a cluster of multiple coccoid cells. (**E**, **F**) Transmission electron micrographs (TEM) of representative cross-sections of (E) the OTI strain and (F) the SH strain, showing the major intracellular compartments including the nucleus, plastid with stacked thylakoid membranes, and storage lipid bodies.

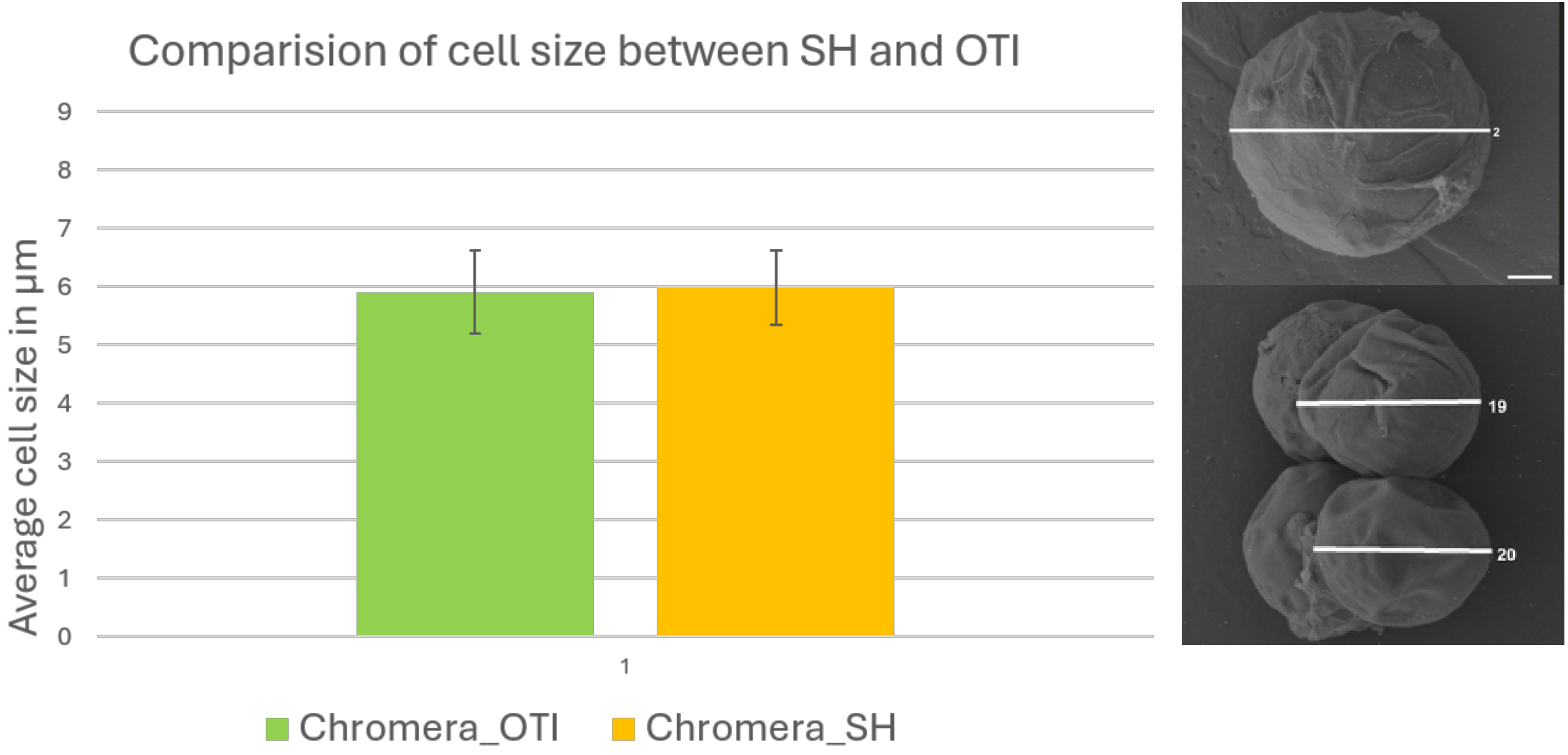

**S2 Fig. Comparison of cell size between the SH and OTI strains of *Chromera velia*.** Bar plot showing the mean maximum cell diameter (Dmax, µm) of the SH (yellow) and OTI (green) strains, quantified from scanning electron micrographs. Individual cell measurements are provided in Supplementary Table 1 (SH: n = 23 cells, mean = 5.98 ± 0.64 µm; OTI: n = 21 cells, mean = 5.91 ± 0.72 µm); error bars represent standard deviation across cells. Cell size did not differ meaningfully between the two strains. Representative micrographs (right) illustrate the coccoid morphology and cells scored for Dmax; white lines indicate the axis along which the maximum diameter was measured, and numbers correspond to cell IDs in Supplementary Table 1

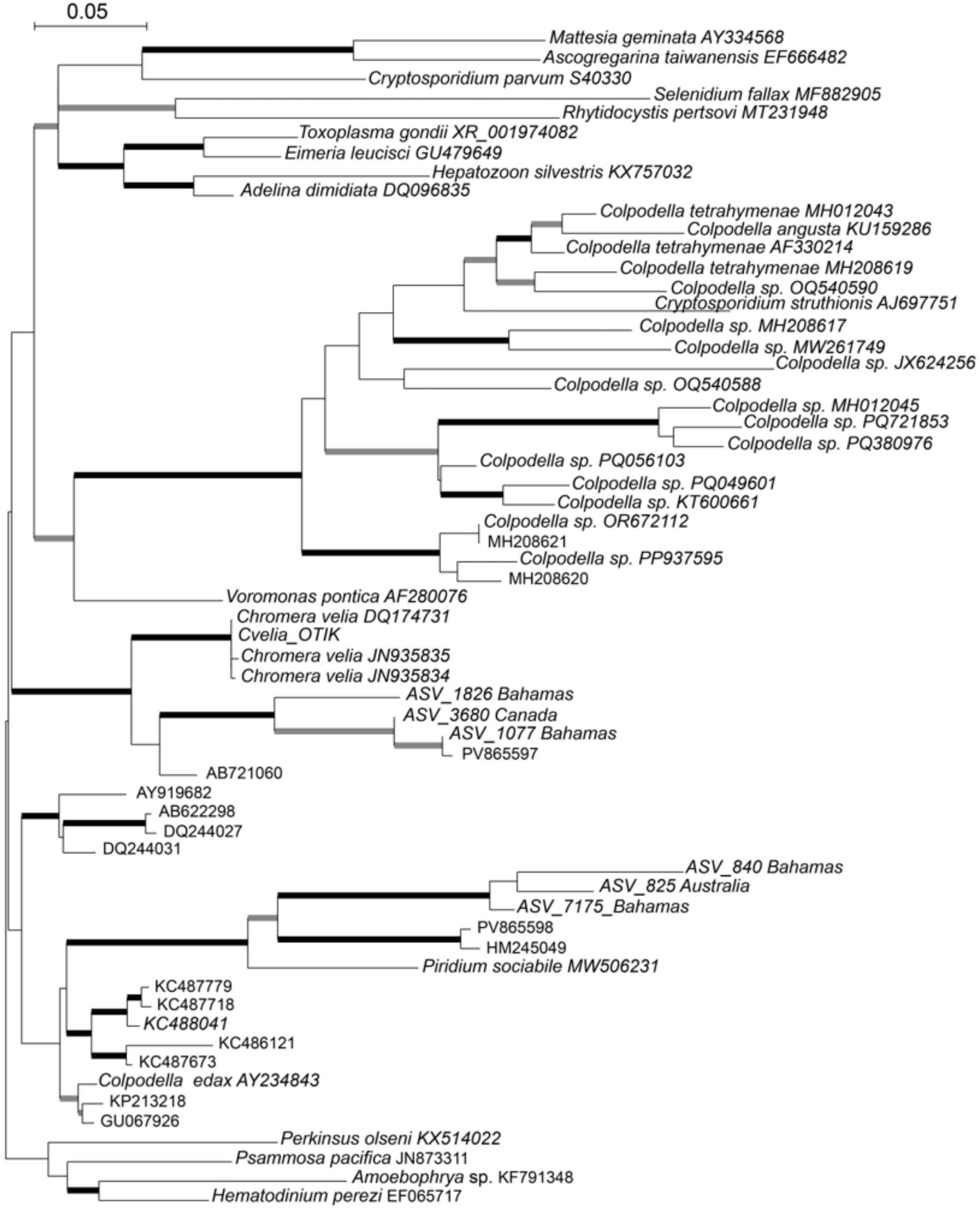

**S3 Fig.** 18S rDNA phylogenetic tree placing the OTI isolate among Apicomplexa.

**S1 Table.** Isolates of *Chromera velia*

| Species | Location | Associated Coral | Isolation temp (° C) | Reference |
|---|---|---|---|---|
| *Chromera velia* (SH) | Sydney harbor, Australia. | Scleractinian coral, Plesiastrea versipora | 20 | [10] |
| Chromera velia (OTI) | One Tree Island on the Great Barrier Reef (GBR), Australia. | Scleractinian coral, Leptastrea purpurea | 26 | [19] |
| Chromera velia | Nelly Bay, Magnetic Island, the inner central region of GBR. | Scleractinian corals, Montipora digitata | 26 | [14,20] |
| Chromera sp. | Caribbean region | Lettuce coral, Agaricia agaricites | - | [21] |

| Chromera_SH measurements | | | Chromera_OT measurements | |
|---|---|---|---|---|
| Cell number | Dmax | | Cell number | Dmax |
| 1 | 6.561 | | 1 | 5.223 |
| 2 | 5.903 | | 2 | 6.706 |
| 3 | 6.101 | | 3 | 6.106 |
| 4 | 4.957 | | 4 | 5.412 |
| 5 | 6.586 | | 5 | 6.248 |
| 6 | 4.861 | | 6 | 4.659 |
| 7 | 5.721 | | 7 | 6.102 |
| 8 | 5.964 | | 8 | 4.953 |
| 9 | 5.756 | | 9 | 5.328 |
| 10 | 7.003 | | 10 | 6.53 |
| 11 | 5.997 | | 11 | 6.833 |
| 12 | 5.838 | | 12 | 5.783 |
| 13 | 5.843 | | 13 | 6.081 |
| 14 | 5.687 | | 14 | 5.687 |
| 15 | 5.165 | | 15 | 6.357 |
| 16 | 5.997 | | 16 | 5.019 |
| 17 | 5.102 | | 17 | 4.934 |
| 18 | 5.263 | | 18 | 5.657 |
| 19 | 7.024 | | 19 | 6.843 |
| 20 | 6.328 | | 20 | 6.539 |
| 21 | 6.878 | | 21 | 7.03 |
| 22 | 6.174 | | average | 5.90619 |
| 23 | 6.769 | | Standard_deviation | 0.715004 |
| Average | 5.977304 | | | |
| Standard_deviation | 0.63747 | | | |

**S2 Table.** Individual cell measurements used to build **S2 Fig.** (SH: n = 23 cells, mean = 5.98 ± 0.64 μm; OTI: n = 21 cells, mean = 5.91 ± 0.72 μm);
